# Deficiency of Mesencephalic Astrocyte-derived Neurotrophic Factor Aggravates Acute Pancreatitis in Mice

**DOI:** 10.1101/2025.07.31.667953

**Authors:** Hui Li, Murong Ma, Wen Wen, Mariah R. Leidinger, Di Hu, Zuohui Zhang, Hong Lin, Jia Luo

## Abstract

Acute pancreatitis (AP) is a complex and potentially severe inflammatory disorder of the pancreas and is one of the most common causes of gastrointestinal hospitalization. Although environmental risk factors such as alcohol and gallstones are well recognized, only a subset of exposed individuals develop AP, suggesting that intrinsic factors, including genetic susceptibility, influence disease onset and progression. Endoplasmic reticulum (ER) stress has emerged as a key mechanism in AP pathogenesis, because ER is essential for protein synthesis, folding, degradation and secretion (proteostasis). Mesencephalic astrocyte-derived neurotrophic factor (MANF), an ER stress-inducible protein highly expressed in the pancreas, plays a critical role in maintaining proteostasis, yet its involvement in AP remains unclear. To investigate the functional role of MANF in AP, we generated pancreas-specific MANF knockout (MANF-KO) mice using the Cre/loxP system and subjected them to moderate experimental AP, that is caerulein- or alcohol-induced AP in mice. In the caerulein model, MANF deficiency exacerbated pancreatic injury in both sexes, as indicated by increased apoptosis (cleaved caspase-3, caspase-12), ER stress markers (eIF2α, p-eIF2α, GRP78), inflammation (IL-6, TNFα), regenerative activity (Ki67), and pancreatic lipase levels. Notably, male MANF-KO mice exhibited enhanced inflammation (HMGB1), macrophage infiltration (CD68), and oxidative stress (DNP, HNE), which were not observed in females. In the alcoholic AP model, both male and female MANF-KO mice showed increased ER stress (p-IRE1, p-eIF2α, GRP78), apoptosis, inflammation, macrophage infiltration, regeneration, and lipase levels, whereas elevated HMGB1 expression and oxidative stress again predominantly occurred in male MANF-KO mice. Together, these findings reveal a critical and sex-specific role for MANF in regulating pancreatic stress responses and inflammatory injury, supporting its potential contribution as a genetic factor in AP pathogenesis.

## Introduction

Acute pancreatitis (AP) is a common inflammatory condition of the pancreas that is often self-limiting and resolves without major medical intervention ^1^. However, severe cases can lead to hospitalization and long-term complications, including recurrent AP, chronic pancreatitis, and other pancreatic disorders such as diabetes and pancreatic cancer ^1–4^.

The development of AP is influenced by both genetic and environmental factors, with gallstones and alcohol consumption being the leading causes ^1, 5^. The incidence and etiology of AP vary by sex: gallstone-related AP is more frequent in women, whereas alcohol-induced AP is more common in men ^6^. Notably, men are more likely to experience worse clinical outcomes, including higher recurrence rates and progression to chronic disease ^2, 7^. Both gallstone- and alcohol-related AP are associated with greater morbidity in men, suggesting potential sex-based differences in disease severity and recovery ^2, 8, 9^. Due to the heterogeneity of clinical presentations and unpredictable disease course, a key challenge in AP management is to identify the risk factors that affect susceptibility, severity and long-term outcomes.

Proteostasis refers to the cellular processes that control the synthesis, folding, trafficking, and degradation of proteins to maintain a functional proteome. Maintaining proteostasis is especially critical in secretory cells-like pancreatic acinar cells-which produce large amounts of digestive enzymes and are therefore under constant protein production stress ^10^. Disruption of proteostasis has been implicated in the pathogenesis of pancreatitis ^11^. The endoplasmic reticulum (ER) is an organelle involved protein synthesis and folding, lipid synthesis, and calcium storage. The ER plays a central role in proteostasis through protein folding, ER-associated degradation (ERAD), and unfolded protein response (UPR) ^12^. ER stress which is induced by excessive accumulation of unfolded/misfolded proteins causes disruption of proteostasis ^12^. Genetic mutations in proteins involved in proteostasis (e.g., PRSS1, SPINK1) increase pancreatitis risk ^13^. Chemical inducers of ER stress (e.g., tunicamycin) can provoke pancreatitis in experimental systems ^14^.

Mesencephalic astrocyte-derived neurotrophic factor (MANF) is an ER-resident protein that is widely expressed across human and mouse tissues, with particularly high levels in secretory organs such as the pancreas ^15–17^. MANF plays a vital role in maintaining ER homeostasis and is essential for normal pancreatic function. In the exocrine pancreas, aberrant MANF expression in acinar and ductal cells has been implicated in the development of chronic alcoholic pancreatitis ^18^. An *in vitro* study using mouse pancreatic acinar cells shows that MANF deficiency exacerbates alcohol-induced ER stress and promotes cell death, whereas exogenous MANF administration mitigates these effects *in vitro* ^19^. Beyond its role in the exocrine compartment, MANF also supports β-cell survival in the endocrine pancreas and has been shown to confer protection in mouse models of diabetes, underscoring its broader importance in pancreatic health ^20, 21^.

In this study, we investigated the role of MANF in AP development using pancreas-specific MANF knockout mice (*Pdx1^Cre/+^*:: *Manf^fl/fl^*; MANF-KO). Utilizing two experimental AP models, the caerulein- and alcohol-induced AP, both of which recapitulate key pathophysiological features of the disease, we examined key cellular processes associated with AP, including ER stress, cell death, inflammatory signaling, macrophage infiltration, oxidative stress, regulation of pancreatic digestive enzymes, and cell proliferation. Our findings indicate that MANF deficiency exacerbates AP development and provide a new insight into the molecular mechanisms by which MANF, as a potential *novel* genetic factor, contributes to the pathogenesis and progression of AP.

## Material and methods

### Materials

Rabbit anti-cleaved caspase-3, rabbit anti-eIF2a, rabbit anti-phospho-eIF2α (Ser51), rabbit anti-Ki67, rabbit anti-Dinitrophenol (DNP), and rabbit anti-CD68 antibodies were obtained from Cell Signaling Technology (Danvers, MA). Rabbit anti-MANF, rabbit anti-phosphor-IRE1 (S724), rabbit anti-IL-6, and rabbit anti-TNFα antibodies were obtained from obtained from AbCam (Cambridge, MA). Rabbit anti-GRP78 was obtained from Novus Biologicals (Littleton, CO). Rabbit Anti-caspase 12 was obtained from Biovision (Mountain View, CA). Rabbit anti-α-amylase antibody was purchased from Proteintech (Rosemont, IL). Rabbit anti-lipase antibodies, amylase activity assay kit and lipase activity assay kit were obtained from Sigma-Aldrich (St. Louis, MO). Goat anti-4-Hydroxynonenal (HNE) antibody was purchased from LifeSpan BioSciences (Seattle, WA). HRP-conjugated anti-rabbit and anti-mouse secondary antibodies were purchased from VWR (Radnor, PA). Anti-goat secondary antibody was obtained from Thermo Fisher (Waltham, MA). Biotin-conjugated anti-rabbit secondary antibody, ABC kit and DAB substrate kit were purchased from Vector Laboratories (Burlingame, CA). Ketamine/xylazine was obtained from Butler Schein Animal Health (Dublin, OH).

### Animal model and treatment

All mice used in this study were bred and housed in the University of Iowa animal facility in accordance with NIH and institutional guidelines. All experimental procedures were approved by the Institutional Animal Care and Use Committee (IACUC) of the University of Iowa. The generation of *Manf^fl/fl^* mice has been described previously ^20, 22^. To generate embryonic deletion of pancreatic MANF knockout mice (*Pdx1^Cre/+^*:: *Manf^fl/fl^*mice), *Pdx-1Cre^6TUV^* transgenic mice (Stock 014647, C57BL/6J background; The Jackson Laboratory) ^23^ were crossed with *Manf^fl/fl^* mice. Littermate *Manf^fl/fl^* mice were used as wild-type (WT) controls. All animals were maintained in a 12-hour/12-hour light/dark cycle at a controlled temperature of 22±1°C with free access to standard chow and water. For the caerulein-induced AP model, 8-week-old mice were received intraperitoneal injections of caerulein (50 μg/kg body weight) every hour for six consecutive hours, as described previously ^24, 25^ (Fig. 1). Control mice received equivalent volumes of PBS under the same conditions. One hour after the final injection, mice were euthanized, and pancreatic tissues and blood samples were collected for histological and biochemical analyses. For the alcohol-induced AP model, 8-week-old mice were administered 25% (w/v) ethanol solution via oral gavage at a dose of 5 g/kg/day for 5 consecutive days (Fig. 1). Control mice received equivalent volumes of water. Four hours after the final gavage, mice were sacrificed, and pancreatic tissues and blood samples were collected for analysis.

**Figure 1.**
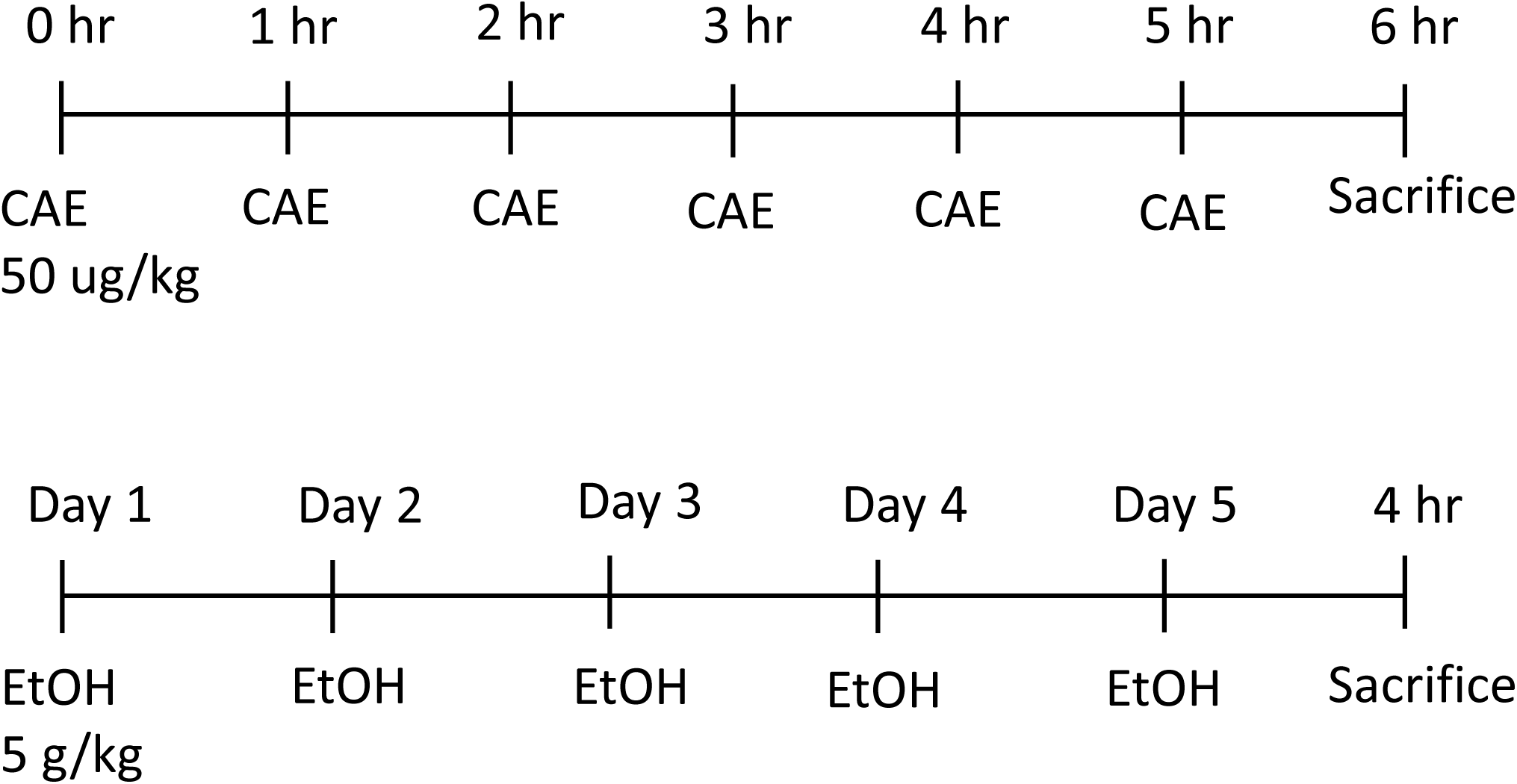
Experimental design for caerulein (CAE)- or alcohol (EtOH)-induced acute pancreatitis (AP) models in mice. In the CAE-induced AP model, 8-week-old *Pdx1^Cre/+^*:: *Manf^fl/fl^* (MANF-KO) and *Manf^fl/fl^*littermates (wild-type, WT) mice as controls received hourly intraperitoneal (i.p.) injections of caerulein (50 µg/kg) for 6 hours. Mice were sacrificed 1 hour after the final injection, and pancreatic tissues and blood were collected. In the EtOH-induced AP model, mice were administered ethanol (5 g/kg, 25% w/v) via oral gavage once daily for 5 consecutive days. Mice were sacrificed 4 hours after the final gavage for tissue and blood collection.

### Dual-energy X-ray absorptiometry (DEXA)

Body composition of mice was assessed using the iNSiGHT DXA System (Scintica, TX) located in the Small Animal Imaging Core (SAIC) facility at the University of Iowa. This non-invasive imaging technique requires the animals to remain still during scanning; therefore, each mouse was briefly anesthetized with inhaled isoflurane. DXA scans were completed in under 5 minutes per animal. The scanning platform was thoroughly disinfected between each session to ensure proper hygiene and prevent cross-contamination.

### Behavioral tests

A battery of behavioral tests was conducted to assess anxiety-like behavior, motor function, and social interaction in mice. The number of animals per experimental group was as follows: for males: WT (n = 9), MANF-KO (n = 8); for females: WT (n = 14), MANF-KO (n = 8). Anxiety-like behaviors were assessed using the open field and elevated plus maze tests. Motor coordination and balance were evaluated using the rotarod test. Social behavior was examined with the three-chamber sociability task. The procedures for these behavioral tests have been previously described ^26^. All behavioral experiments were conducted at the Neural Circuits and Behavior Core (NCBC) at the University of Iowa when mice were 8 weeks old. All animals were habituated to the testing room for at least 30 minutes prior to each behavioral assessment. To minimize potential carryover effects, animals were given at least one day of rest between tests. The testing schedule was as follows: open field (Days 1–2), elevated plus maze (Day 5), three-chamber sociability (Day 8), and rotarod (Day 10).

#### Open field

Each mouse was placed in the center of a custom-built open-field chamber (16″ × 16.5″ × 12″) featuring opaque black walls and a white floor. The central zone of the arena was defined as the area occupying the middle half of the chamber’s length and width. Behavioral testing was conducted under uniform lighting conditions of 110–130 lux. Each animal’s activity was recorded for 10 minutes per session, once daily for two consecutive days, to assess anxiety-like behavior and locomotor activity over time. Videos were analyzed using the EthoVision XT 17 tracking system (Noldus Information Technology, Wageningen, Netherlands), and key parameters including total distance traveled and time spent in the center zone were quantified.

#### Elevated plus maze

Anxiety-like behavior was evaluated using the elevated plus maze (San Diego Instruments, San Diego, CA). The maze was illuminated by an 18-inch ring light (Inkeltech) mounted around an overhead camera, producing approximately 215 lux in the open arms. Each animal was placed in the central zone of the maze, and its activity was recorded for 5 minutes. The time spent in and number of entries into the open and closed arms were quantified using the EthoVision XT 17 tracking system.

#### Rotarod

Motor coordination and balance were assessed using an accelerating rotarod apparatus (IITC Life Science, Woodland Hills, CA). During the training session, mice were placed on the rotarod rotating at a constant speed of 4 revolutions per minute (rpm) for 5 minutes to acclimate them to the apparatus. For the testing phase, the rod was programmed to accelerate linearly from 4 to 40 rpm over a 5-minute period. The latency to fall was automatically recorded for each mouse. Mice underwent three test trials, with a rest period of 10–15 minutes between trials. The average latency to fall across the three trials was calculated for each animal.

#### Three-chamber sociability task

The test was conducted in a custom-made, matte black plastic three-chambered arena, with openings allowing free movement between chambers. Each of the two side chambers contained a perforated plastic cylinder, enabling interaction between the test mouse and either an object or another mouse placed inside the cylinder. During the 10-minute habituation session, both cylinders were empty, and the test mouse was placed in the center chamber and allowed to explore the arena freely. This was followed by a 10-minute testing session, during which a novel object was placed in one cylinder and an unfamiliar, age-matched (4–5 months old), same-sex wild-type C57BL/6 mouse was placed in the other. The test mouse had no prior exposure to the novel mouse. Time spent and number of entries into each chamber, as well as direct interactions with the cylinders, were recorded and analyzed using EthoVision XT 17 software.

### Tissue preparation and immunoblotting

Mice were anesthetized via intraperitoneal injection of ketamine (100 mg/kg) and xylazine (10 mg/kg). The pancreas was dissected, snap-frozen in dry ice, and stored at −80°C. Protein extraction and immunoblotting (IB) were performed as previously described ^27, 28^. Briefly, pancreatic tissues were homogenized in ice-cold 1× RIPA buffer (diluted from 10×; Cell Signaling Technology) containing 20 mM Tris-HCl (pH 7.5), 150 mM NaCl, 1 mM Na₂EDTA, 1 mM EGTA, 1% NP-40, 1% sodium deoxycholate, 2.5 mM sodium pyrophosphate, 1 mM β-glycerophosphate, 1 mM Na₃VO₄, 1 µg/mL leupeptin, and 1 mM PMSF. Homogenates were centrifuged at 13,000 × g for 15 minutes at 4°C, and the supernatants were collected. Protein concentrations were determined, and 40 µg of each sample was separated by SDS-polyacrylamide gel electrophoresis and transferred onto nitrocellulose membranes. Membranes were blocked in 5% BSA in 1× TBST (137 mM NaCl, 20 mM Tris-HCl, pH 7.6, with 0.1% Tween-20) for 1 hour at room temperature, followed by overnight incubation at 4°C with primary antibodies. After three 10-minute washes in TBST, membranes were incubated with HRP-conjugated secondary antibodies for 1 hour at room temperature. Signal detection was performed using enhanced chemiluminescence (GE Healthcare), and band intensities were quantified using Image Lab 6.1 software (Bio-Rad, Hercules, CA).

### Immunohistochemistry

Immunohistochemistry (IHC) was performed as previously described with minor modifications ^29^. Briefly, paraffin-embedded pancreatic tissues were sectioned at 5 μm thickness using a microtome and mounted onto positively charged slides. Sections were deparaffinized, rehydrated, and subjected to antigen retrieval as needed. Non-specific binding was blocked with a solution containing 1% BSA, 2% goat serum, and 0.1% Triton X-100 for 1 hour at room temperature. Slides were then incubated overnight at 4°C with primary antibodies against cleaved caspase-3 (1:200), MANF (1:400), TNFα (1:100), HMGB (1:400), CD68 (1:200), or Ki-67 (1:400). Negative controls were included by omitting the primary antibody. After washing with PBS, sections were incubated with biotinylated goat anti-rabbit IgG (1:200) for 1 hour at room temperature, followed by incubation with the avidin–biotin–peroxidase complex (1:100 in PBS) for another hour. Immunoreactivity was visualized using 0.05% 3,3′-diaminobenzidine (DAB) containing 0.003% H₂O₂ in PBS. Positive cells were quantified under ×40 magnification by counting five randomly selected microscopic fields from four mice per group.

### Measurement of activity of plasma amylase and lipase

Plasma samples were isolated and stored at −80 °C until enzymatic activity assays were performed. The activities of α-amylase and lipase were assessed using commercial assay kits (Amylase Activity Assay Kit and Lipase Activity Assay Kit; Sigma-Aldrich, St. Louis, MO) according to the manufacturer’s instructions. For each assay, 2 μL of plasma was diluted to a final volume of 50 μL. Enzyme activities were quantified and reported as units per liter (U/L), where one unit corresponds to the formation of 1 nmol of product per minute per milliliter (nmol/min/mL).

### Data Analysis

A two-way analysis of variance (ANOVA) was performed to assess the effects of sex and treatment on the relative abundance of selected proteins. Post hoc comparisons were conducted using Tukey’s multiple comparison test to identify significant differences between groups. All statistical analyses were carried out using GraphPad Prism version 10.5.0 (GraphPad Software, Inc.), with a significance level set at *p* < 0.05.

## Results

### Characterization of pancreatic MANF-KO mice

To investigate the role of MANF in pancreatic function, we generated pancreas-specific MANF knockout (MANF-KO) mice using the Cre/loxP system. *Pdx1^Cre/+^*:: *Manf^flox/+(fl/+)^* mice were crossed with *Manf^flox/flox(fl/fl)^* mice to produce *Pdx1^Cre/+^*:: *Manf^fl/fl^* offspring (hereafter referred to as MANF-KO), in which MANF is constitutively deleted in the pancreas. The Pdx1-Cre driver, under the control of the pancreatic and duodenal homeobox 1 (Pdx1) promoter, targets all pancreatic cell types, including both endocrine islets and exocrine acinar cells ^30^. In this study, we focused on the exocrine pancreas and used *Manf^fl/fl^* littermates as wild-type (WT) controls. Successful pancreatic deletion of MANF was confirmed by immunoblotting and immunohistochemistry (Figs. 2B and 2C). MANF-KO mice exhibited notable phenotypic differences, including reduced body weight and pancreatic mass (Fig. 2A), as well as increased urination. Dual-energy X-ray absorptiometry (DEXA) analysis further revealed lower body fat percentages and reduced femoral bone mineral density (BMD), particularly in female MANF-KO mice (Figs. 2D–2G). To determine whether MANF KO affects neurobehaviors, we evaluated anxiety-like behaviors, motor coordination and sociability in these mice. Pancreatic MANF deficiency had a modest effect on behavioral outcomes. Male MANF-KO mice exhibited increased anxiety-like behavior on day 1 of a two-day open field test, while female MANF-KO mice displayed anxiolytic behavior on day 2 (Supplementary Fig. 1). MANF deficiency did not affect other behaviors tested, namely, motor coordination and sociability.

**Figure 2.**
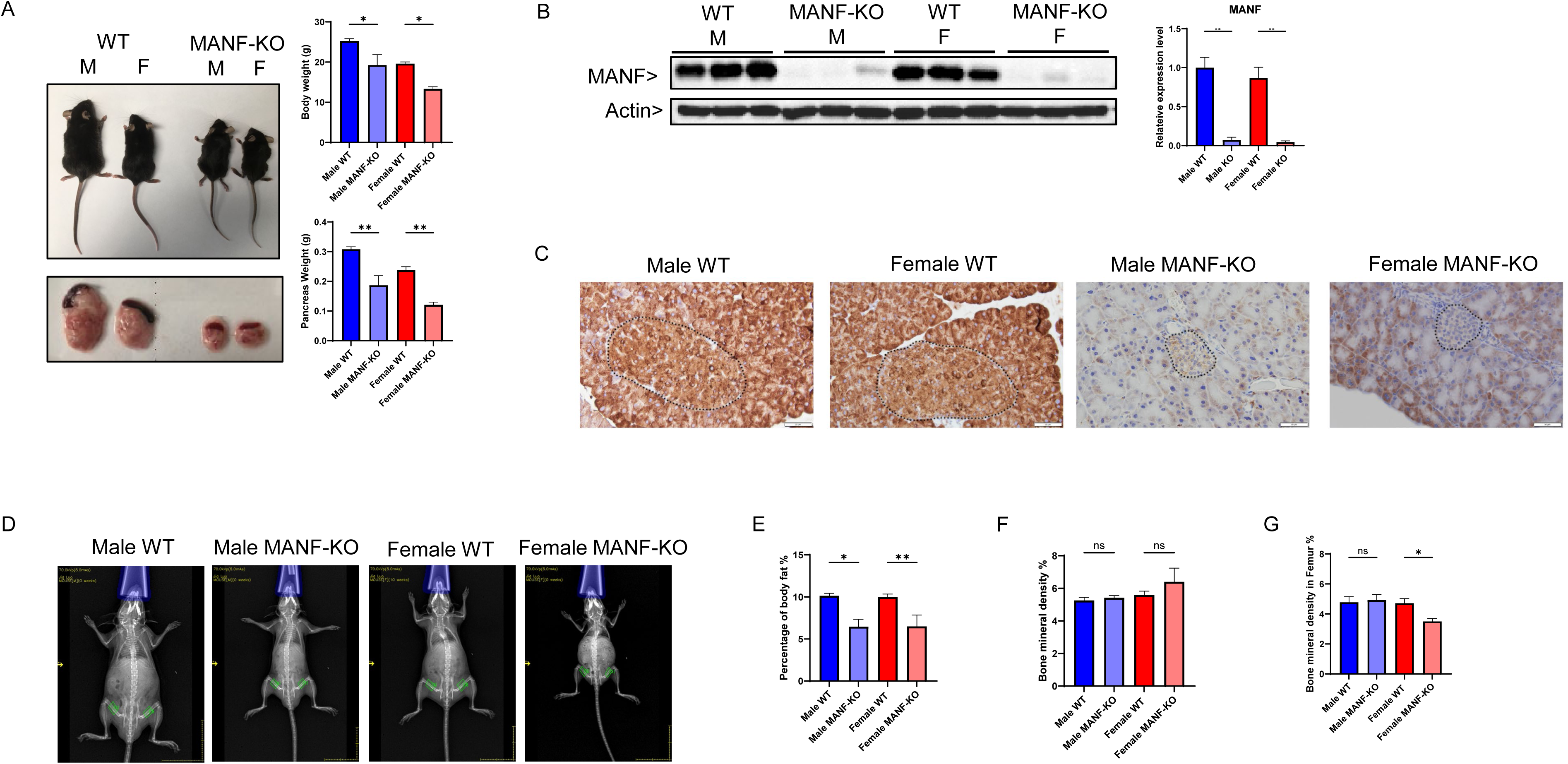
Characterization of *Pdx1^Cre/+^*:: *Manf^fl/fl^* (MANF-KO) mice. **A**. Body weight (top panel) and pancreatic mass (bottom panel) of MANF-KO mice and their *Manf^fl/fl^* (wild-type, WT) littermates were measured at 8 weeks of age in both males and females. n = 5 for male WT, n = 5 for male KO, n = 5 for female WT, n = 7 for female KO. **B**. A representative immunoblot (IB) image showing the expression of MANF in pancreatic tissue (left) and the densitometric quantification (right). The levels of MANF expression were normalized to the expression of actin. n = 3 per group. **C**. Representative images of immunohistochemistry (IHC) staining of MANF in pancreatic tissues, counterstained with hematoxylin (magnification of 40X). The pancreatic islets are indicated by dotted outlines. **D–G**: Assessment of body composition and bone density using dual-energy X-ray absorptiometry (DEXA): D. Representative images of DEXA scans; E. quantification of body fat percentage; F. total bone mineral density (BMD), and G. femoral BMD. n = 5 for male WT, 4 for male KO, 6 for female WT, and 5 for female KO. Statistical comparisons were performed using two-way ANOVA followed by Tukey’s post hoc test. Data are presented as mean ± SEM. Significance is indicated as follows: *p < 0.05, **p < 0.01, ***p < 0.001, ****p < 0.0001; ns = not significant (p > 0.05).

### Effects of MANF deficiency on apoptosis in the pancreas of caerulein- and alcohol-induced AP models

To explore the role of MANF in pancreatic cell death during acute AP, we utilized both caerulein- and alcohol-induced AP models in 8-week-old male and female *Manf^fl/fl^* (WT) and *Pdx1^Cre/+^*:: *Manf^fl/fl^* (MANF-KO) mice. In the caerulein-induced model, mice received six hourly intraperitoneal (IP) injections of caerulein (50 μg/kg body weight), and pancreatic tissues were collected one hour after the final injection. In the alcohol-induced model, mice were administered ethanol (25% w/v, 5 g/kg/day) via oral gavage for five consecutive days and sacrificed six hours after the final dose (Fig. 1). Apoptotic activity was assessed by measuring cleaved caspase-3 (c-cas-3) expression via immunoblotting (IB) and immunohistochemistry (IHC). In WT controls, neither caerulein nor alcohol alone induced significant cell death in either sex. However, MANF deficiency markedly elevated c-cas-3 levels, and it interacted with either caerulein or alcohol exposure resulting in a further increase in apoptosis in both sexes (Figs. 3A and 3B). IHC analysis further revealed that c-cas-3 staining was predominantly localized to pancreatic acinar cells (Fig. 3C and 3D). To evaluate ER stress-associated apoptosis, we examined caspase-12 expression. Neither caerulein nor alcohol treatment alone, nor MANF deficiency by itself, significantly altered caspase-12 levels. However, the combination of MANF deletion with either caerulein or ethanol treatment significantly increased caspase-12 expression in both sexes, indicating enhanced ER stress-mediated apoptosis (Figs. 3A and 3B). Histological analysis of H&E-stained pancreatic sections showed that caerulein or alcohol alone induced only mild AP features, including acinar cell vacuolation, intercellular edema, and limited structural disruption (Fig. 3E). In contrast, MANF-deficient mice treated with either agent exhibited severer AP pathology characterized by widespread edema, intense inflammatory infiltration, and extensive acinar cell necrosis (Fig. 3E).

**Figure 3.**
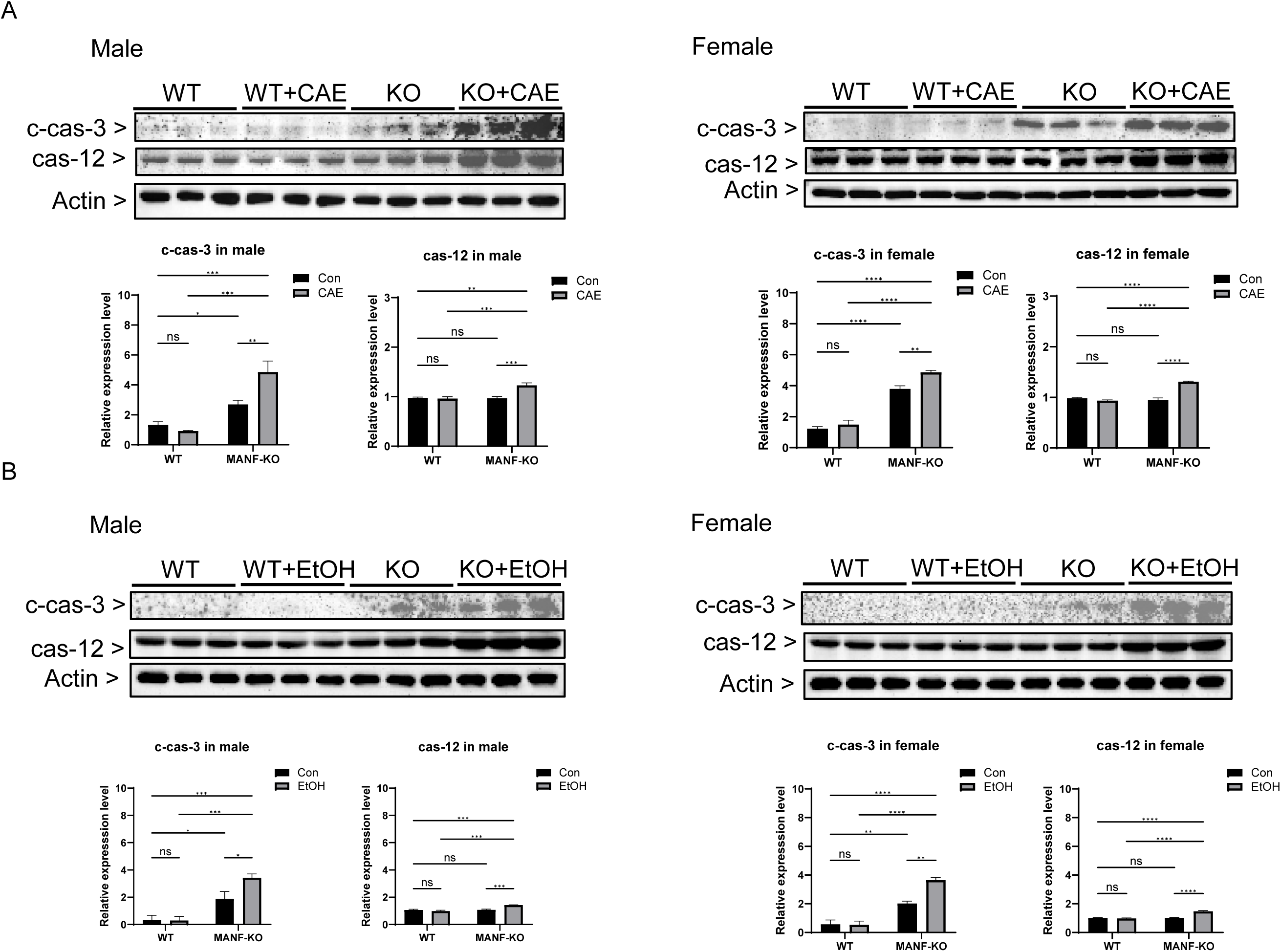
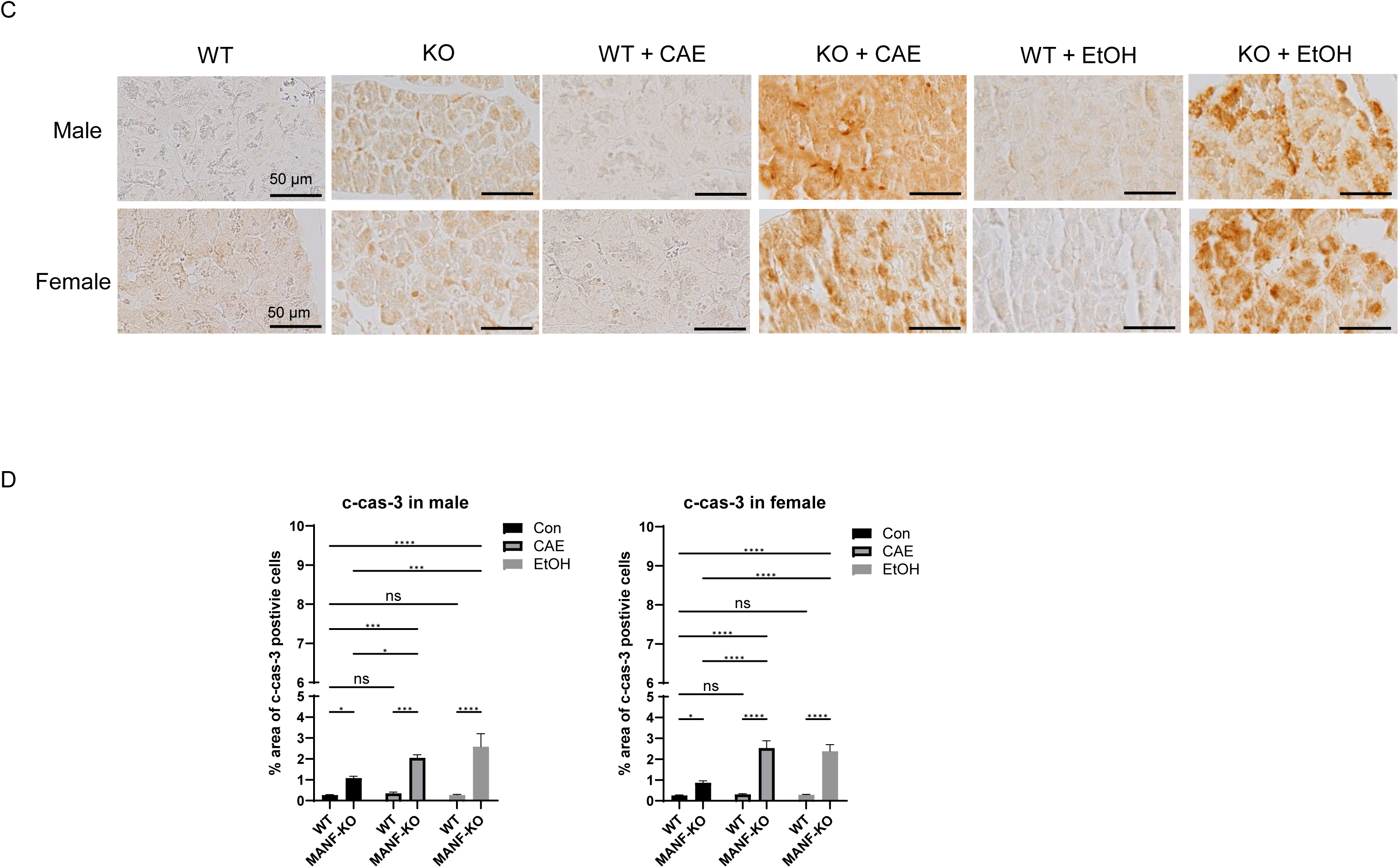
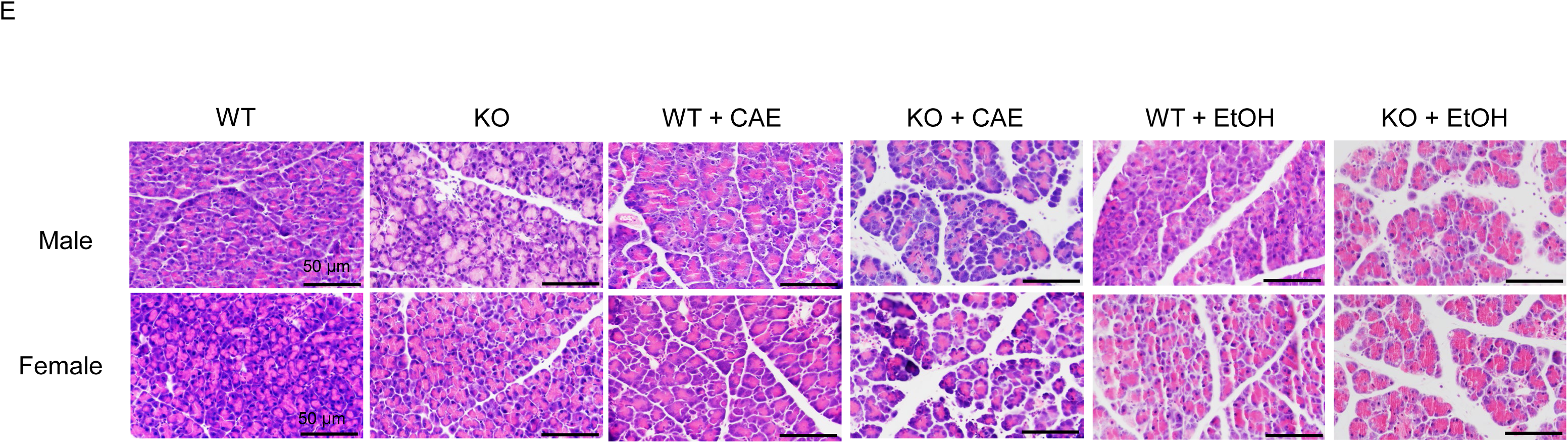
Effects of MANF deficiency on apoptosis in the pancreas of caerulein- and alcohol-treated mice. **A**. Male (left) and female (right) *Pdx1^Cre/+^*:: *Manf^fl/fl^* (MANF-KO) and *Manf^fl/fl^* (wild-type, WT) mice were treated with caerulein as described in Fig. 1. The expression of cleaved caspase-3 (c-cas-3) and caspase-12 (cas-12) in pancreatic tissues was evaluated by IB analysis and their expression levels were quantified and normalized to actin levels. n = 3 per group. **B**. The mice were exposed to alcohol as described in Fig. 1. The expression of c-cas-3 and cas-12 in pancreatic tissues was evaluated and quantified by IB analysis and their expression levels were normalized to actin levels. n = 3 per group. **C.** Representative immunohistochemistry (IHC) images of c-cas-3 staining (magnification of 40X) in pancreatic tissues. **D.** Quantification of c-cas-3-positive cells from five randomly selected fields within the pancreas of each mouse. n = 4 mice per group. **E**. Representative images of hematoxylin and eosin (H&E) staining (magnification of 40X) illustrating pancreatic histopathology. Quantitative data are presented as mean ± SEM. Statistical analysis was performed using two-way ANOVA followed by Tukey’s post hoc test. Significance levels: *p < 0.05, **p < 0.01, ***p < 0.001, ****p < 0.0001; ns = not significant (p > 0.05).

### Effects of MANF deficiency on ER stress/UPR in the pancreas of caerulein- or alcohol-induced AP models

To investigate the interaction of MANF and ER stress/UPR pathways in AP, we performed an IB analysis to evaluate the expression of key UPR components, including glucose-regulated protein 78 (GRP78), phosphorylated inositol-requiring enzyme 1 (p-IRE1), eukaryotic translation initiation factor 2A (eIF2A), and its phosphorylated form (p-eIF2A), alongside MANF. In the caerulein-induced AP model, neither caerulein treatment nor MANF deficiency alone altered the expression of these ER stress markers. However, their combination significantly increased the expression of GRP78, eIF2A, and p-eIF2A in both male and female mice (Fig. 4A). Similarly, in the alcohol-induced AP model, ER stress marker levels remained unchanged with either alcohol exposure or MANF deletion alone. In contrast, the combined treatment markedly upregulated p-IRE1, GRP78, and p-eIF2A expression in both sexes, indicating that MANF deficiency promoted ER stress/UPR activation (Fig. 4B).

**Figure 4.**
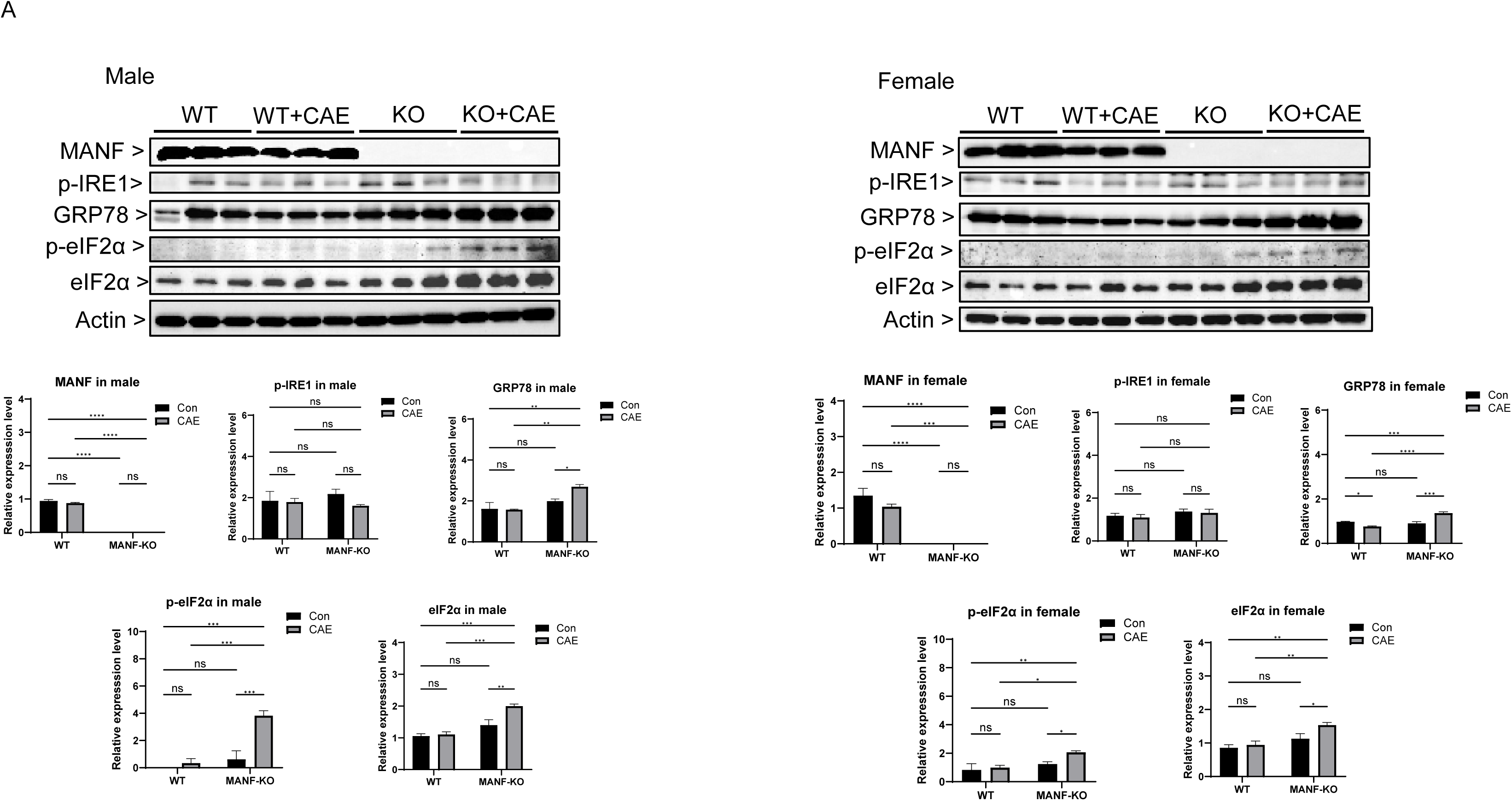
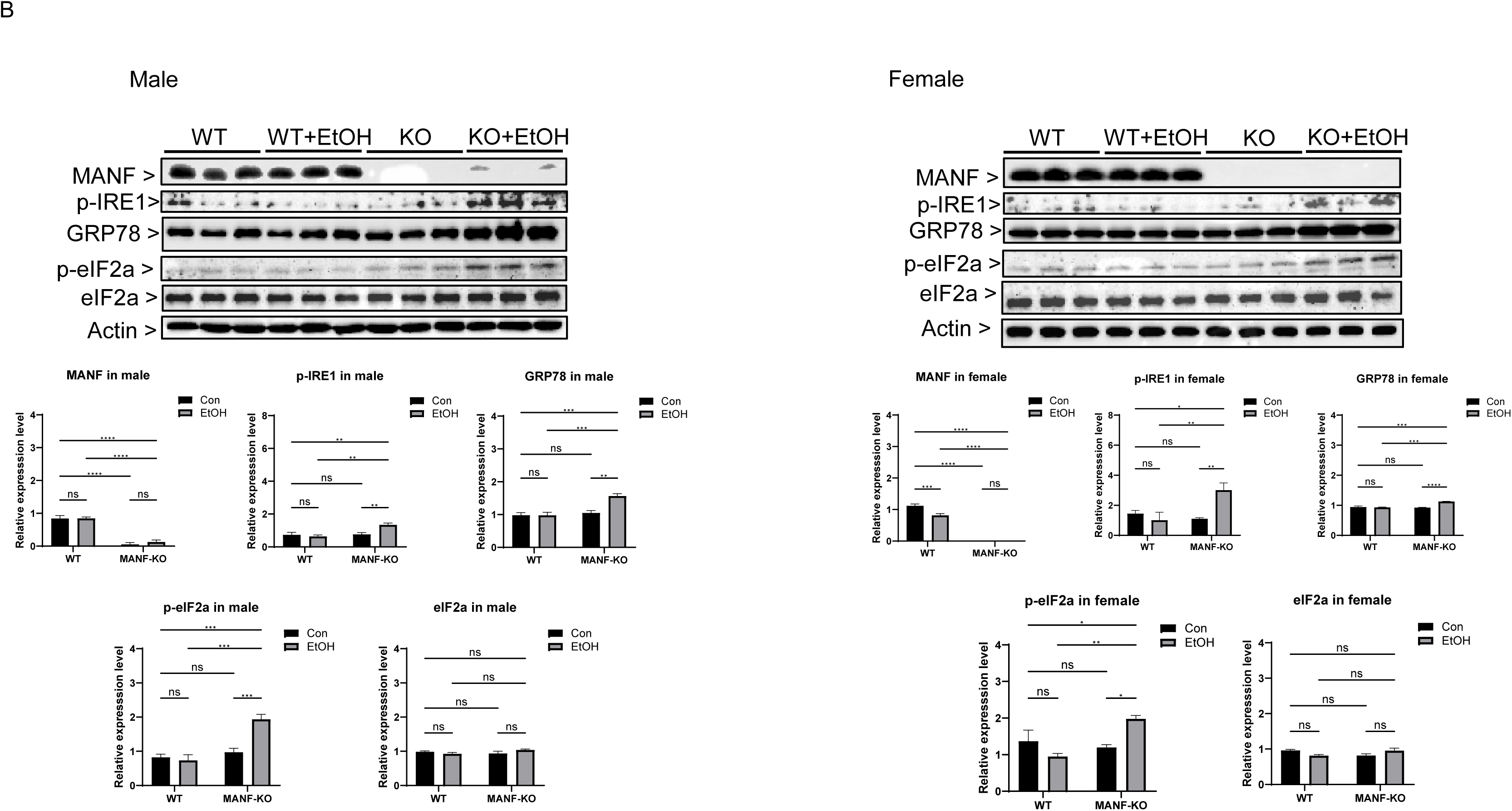
Effects of MANF deficiency on ER stress in the pancreas of caerulein- and alcohol-treated mice. **A**. Male (left) and female (right) MANF-KO and WT mice were treated with caerulein as described in Fig. 1. The expression of phosphorylated IRE1 (p-IRE1), GRP78, total eIF2α, and phosphorylated eIF2α (p-eIF2α) in pancreatic tissues was examined by IB analysis. **B**. Male (left) and female (right) MANF-KO and WT mice were exposed to alcohol as described in Fig. 1 and the expression of UPR proteins was analyzed by IB. n = 3 per group. Data are presented as mean ± SEM and were analyzed by two-way ANOVA followed by Tukey’s post hoc test. Statistical significance: *p < 0.05, **p < 0.01, ***p < 0.001, ****p < 0.0001; ns = not significant (p > 0.05).

### Effects of MANF deficiency on caerulein and alcohol-induced inflammation in the pancreas

We next assessed the inflammatory response in both AP models by evaluating the expression of interleukin-6 (IL6), tumor necrosis factor-alpha (TNFα), and high mobility group box 1 (HMGB1) via immunoblotting (Figs. 5A and 5B). In the caerulein-induced AP model, caerulein alone significantly increased IL6 expression in both sexes but had no effect on TNFα. MANF deficiency independently elevated both IL6 and TNFα levels in males and females, and its combination with caerulein produced a further increase in these proinflammatory cytokines. HMGB1 levels remained unchanged in males with either caerulein or MANF deficiency alone; however, caerulein alone reduced HMGB1 expression in females. Notably, the combination of MANF deficiency and caerulein treatment resulted in a marked increase in HMGB1 expression in males but further suppressed HMGB1 in females. A similar pattern was observed in the alcohol-induced AP model. Alcohol treatment alone selectively increased IL6 levels in males without affecting TNFα in either sex. When combined with MANF deficiency, alcohol significantly elevated both IL6 and TNFα expression in males and females. HMGB1 levels were unaffected by alcohol or MANF deficiency alone, yet their combination significantly elevated HMGB1 expression in males, mirroring the caerulein model, while females showed no such increase. IHC analysis confirmed that TNFα and HMGB1 were predominantly localized to pancreatic acinar cells in both AP models (Figs. 5C, 5D, 5E and 5F).

**Figure 5.**
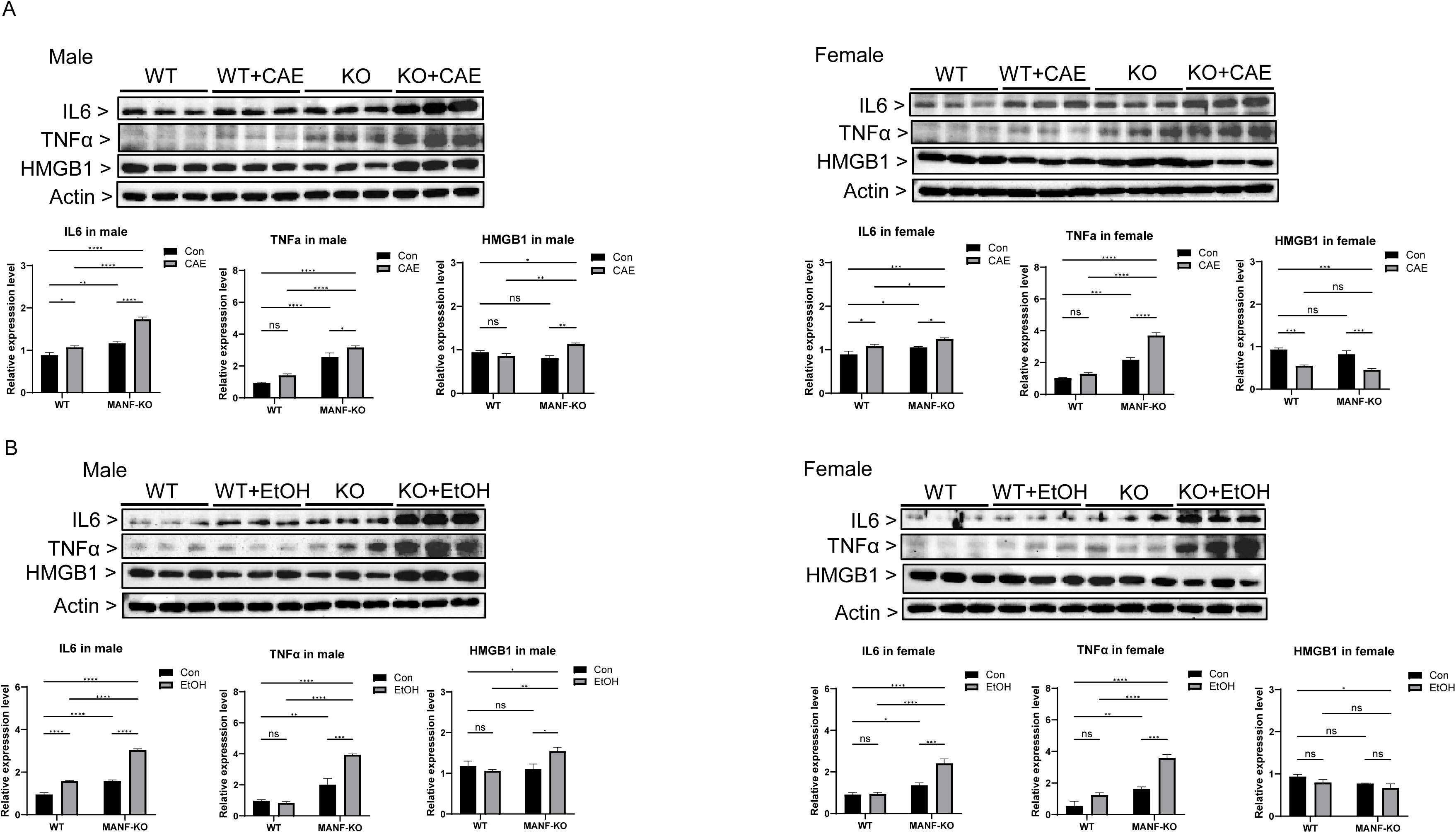
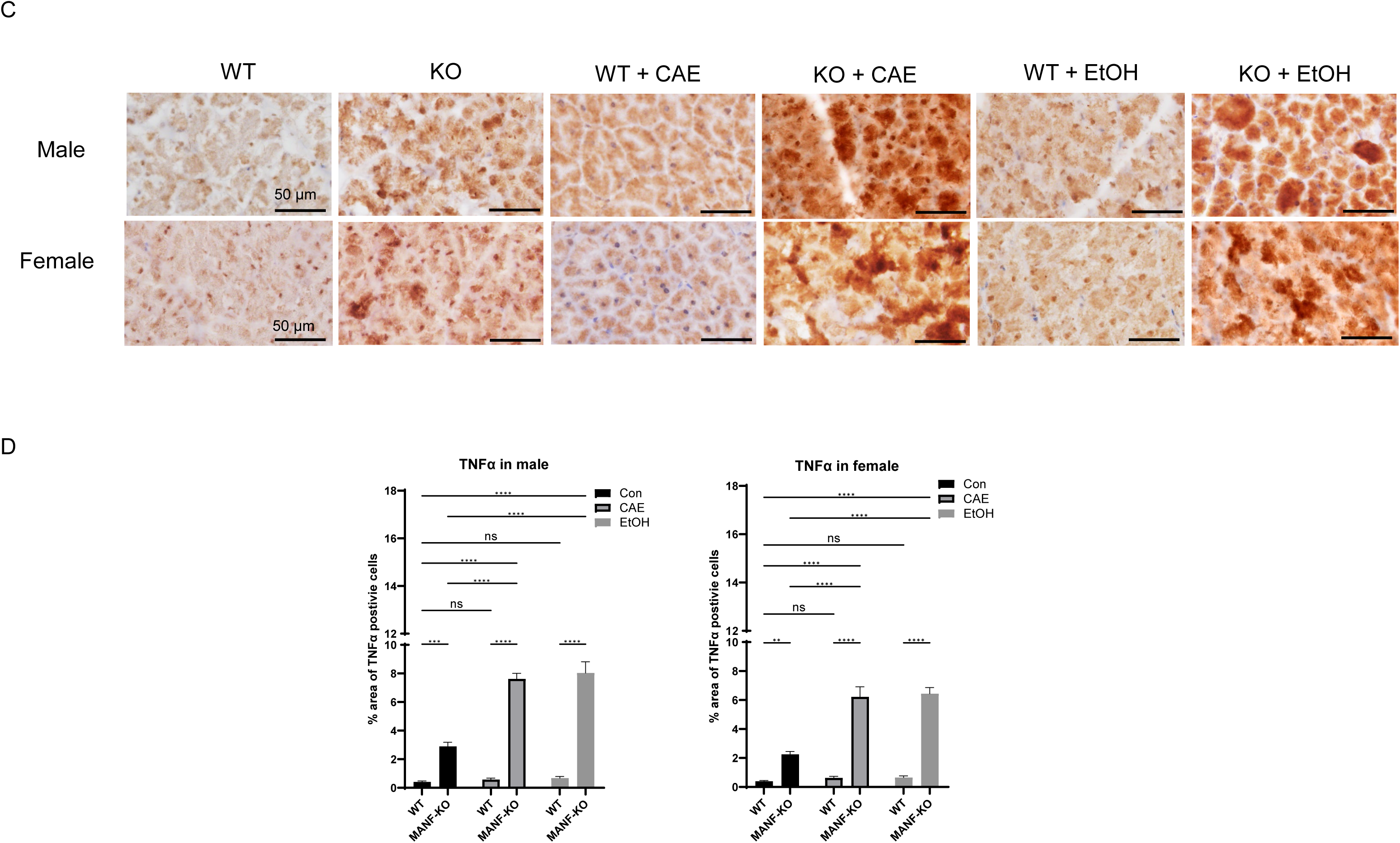
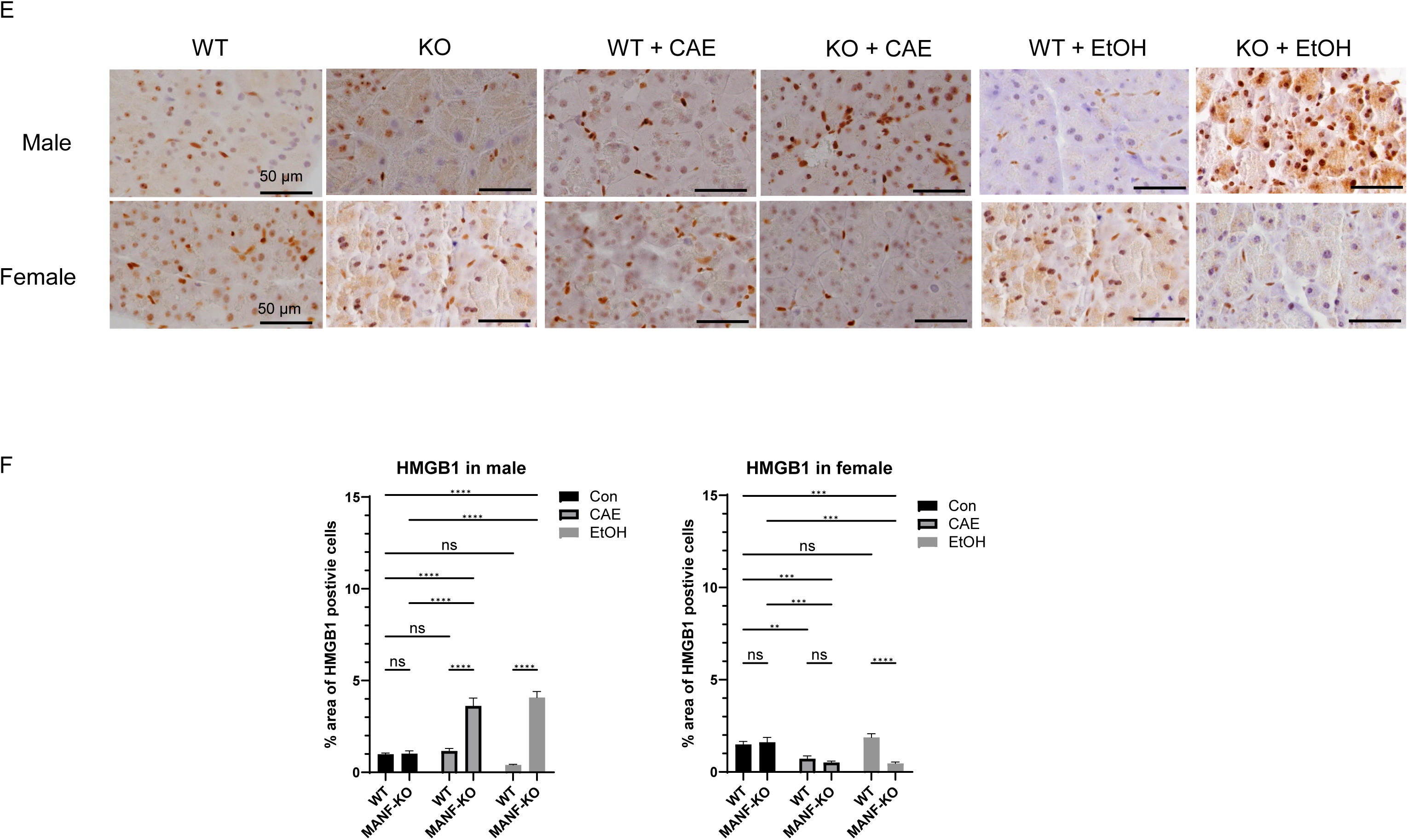
Effects of MANF deficiency on inflammation in the pancreas of caerulein- and alcohol-treated mice. **A.** Male (left) and female (right) MANF-KO and WT mice were treated with caerulein as described above. Representative images of IB showing the expression and quantification of inflammation markers IL6, TNFα and HMGB1 in pancreatic tissues. n = 3 per group. **B**. Male (left) and female (right) MANF-KO and WT mice were exposed to alcohol as described above and the expression of inflammation makers was analyzed by IB. n = 3 per group. **C, E**. Representative images of IHC for TNFα (C) and HMGB1 (E) (magnification of 40X) in pancreatic tissues from caerulein- and alcohol-treated mice. **D, F.** Quantification of TNFα-positive (D) and HMGB1-positive (F) positive cells from five randomly selected fields within the pancreas of each mouse. n = 4 mice per group. Data are presented as mean ± SEM and were analyzed by two-way ANOVA followed by Tukey’s post hoc test. Statistical significance: *p < 0.05, **p < 0.01, ***p < 0.001, ****p < 0.0001; ns = not significant (p > 0.05).

### *Effects of MANF deficiency on macrophage infiltration in* caerulein *and alcoholic AP models*

We also examined the expression of CD68, a macrophage marker indicative of inflammation and oxidative stress responses, in both AP models. In caerulein-induced AP model, CD68 expression was significantly elevated in male mice, whereas no such increase was observed in females (Figs. 6A and 6C). MANF deficiency further increased caerulein-induced CD68 expression in male mice but not female mice (Figs. 6A and 6C). Similarly in alcoholic AP model, alcohol exposure alone increased CD68 levels in male but not female mice. MANF deficiency plus alcohol induced CD68 expression in both sexes (Figs. 6B and 6C). MANF deficiency-stimulated macrophage infiltration was evident by IHC analysis (Fig. 6C and 6D).

**Figure 6.**
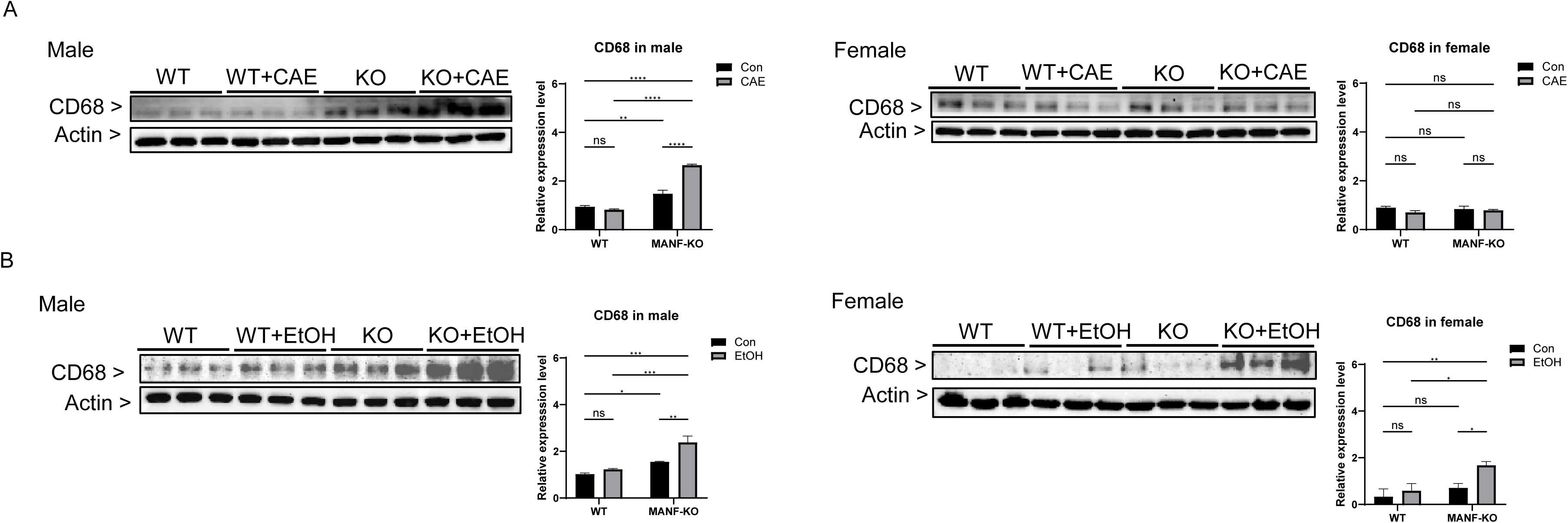
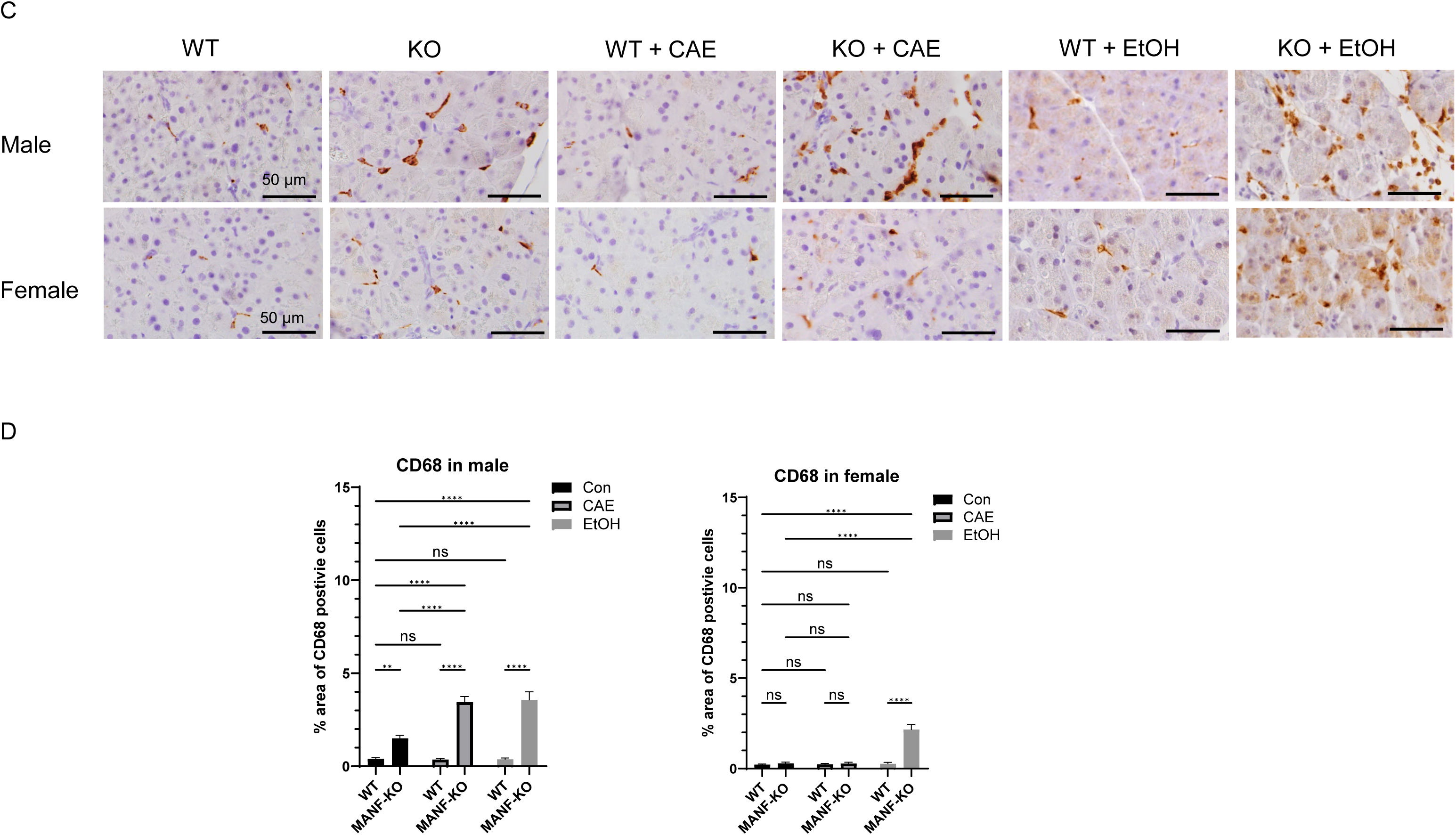
Effects of MANF deficiency on macrophage infiltration in the pancreas of caerulein- and alcohol-treated mice. **A.** Male (left) and female (right) MANF-KO and WT mice were treated with caerulein as described above. Representative images of IB showing the expression and quantification of the macrophage marker CD68 in pancreatic tissues. n = 3 per group. **B.** Male (left) and female (right) MANF-KO and WT mice were exposed to alcohol as described above and the expression of CD68 was analyzed by IB. n = 3 per group. **C**. Representative IHC images (magnification of 40X) showing CD68-positive cells (brown) in pancreatic tissues, counterstained with hematoxylin (blue). **D.** Quantification of CD68-positive cells from five randomly selected fields per pancreas. n = 4 mice per group. Data are presented as mean ± SEM and were analyzed by two-way ANOVA followed by Tukey’s post hoc test. Statistical significance: *p < 0.05, **p < 0.01, ***p < 0.001, ****p < 0.0001; ns = not significant (p > 0.05).

### Effects of MANF deficiency on caerulein- and alcohol-elicited oxidative stress in the pancreas

We further explored the impact of MANF deficiency on oxidative stress by evaluating protein oxidation and lipid peroxidation in both AP models via immunoblotting. Protein oxidation was assessed by measuring protein carbonyl content using IB analysis with anti-dinitrophenol (DNP) antibodies, while lipid peroxidation was evaluated through the detection of 4-hydroxynonenal (4-HNE) adducts ^27, 28^. Our findings revealed sex-specific differences in oxidative stress responses. In males, caerulein treatment alone did not alter 4-HNE levels but significantly reduced DNP levels. MANF deficiency alone had no effect on either marker; however, the combination of MANF deficiency and caerulein treatment led to a marked increase in both HNE and DNP levels, specifically in males (Fig. 7A). Similarly, in alcoholic AP model, neither alcohol nor MANF deficiency alone significantly affected HNE or DNP levels, but their combined treatment resulted in a significant elevation of both markers in males but not females (Fig. 7B).

**Figure 7.**
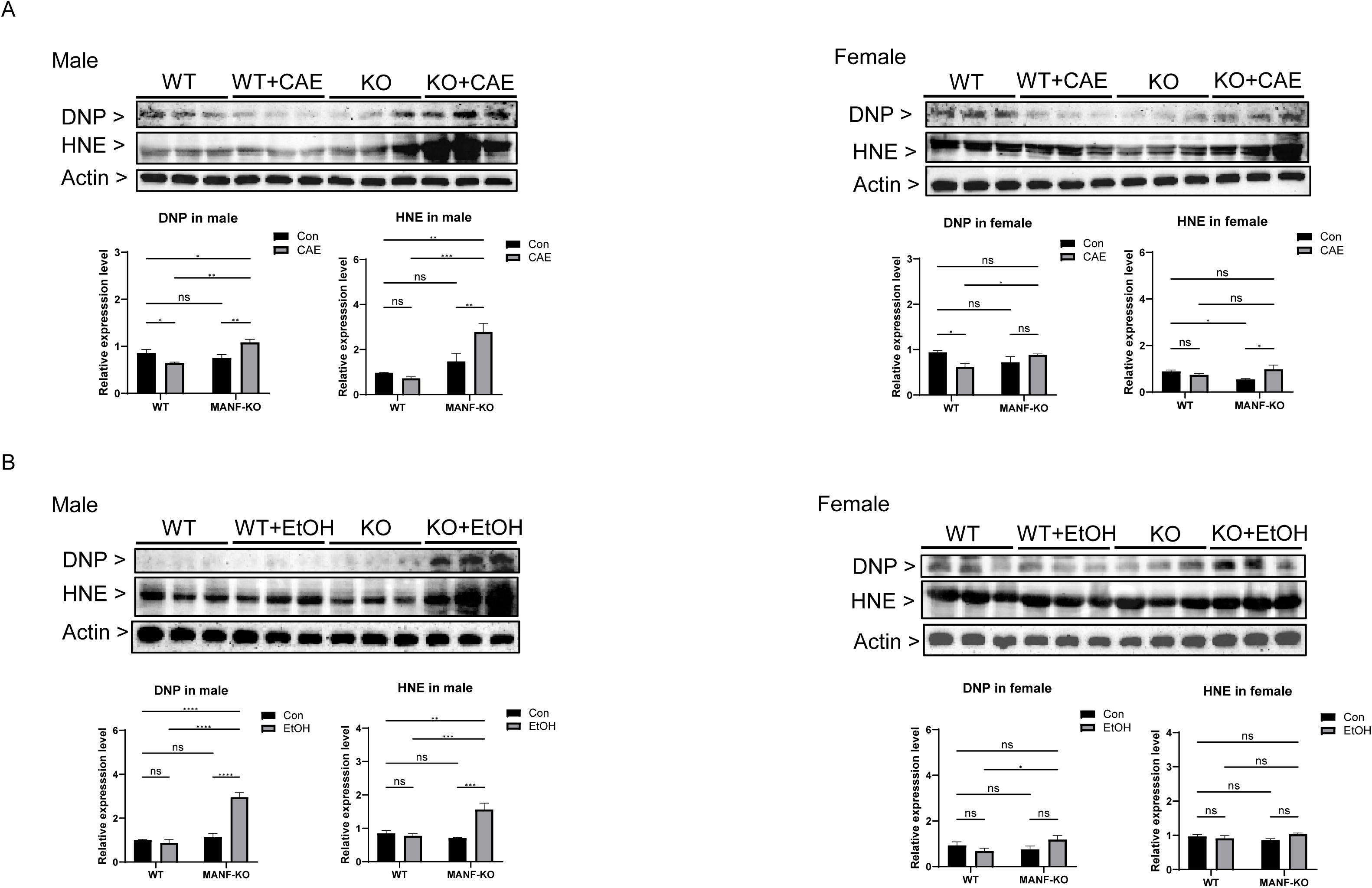
Effects of MANF deficiency on oxidative stress in the pancreas of caerulein- and alcohol-treated mice. **A**. Representative IB images showing the expression of oxidative stress markers, protein carbonylation (DNP) and lipid peroxidation (4-HNE), in pancreatic tissues of male (left) and female (right) mice following caerulein treatment. **B**. Representative IB images showing the expression of DNP and 4-HNE in pancreatic tissues of male (left) and female (right) mice following alcohol exposure. n = 3 per group. Quantified data are expressed as mean ± SEM and analyzed using two-way ANOVA with Tukey’s post hoc test. *p < 0.05, **p < 0.01, ***p < 0.001, ****p < 0.0001; ns = not significant (p > 0.05).

### Effects of MANF deficiency on pancreatic enzymes in caerulein- or alcohol-induced AP models

To assess the impact of MANF deletion on pancreatic digestive enzymes in both AP models, we analyzed amylase and lipase levels in pancreatic tissue and plasma. IB analysis of pancreatic lysates showed that neither caerulein nor alcohol treatment alone altered amylase or lipase expression in males or females (Figs. 8A and 8B). However, MANF deficiency significantly decreased pancreatic amylase levels in both sexes. Interestingly, this reduction was reversed when MANF deletion was combined with either caerulein or alcohol treatment, suggesting a compensatory response under inflammatory conditions. In contrast, MANF deletion markedly increased pancreatic lipase expression in both males and females, an effect that was further amplified when combined with either caerulein or alcohol, indicating a synergistic interaction. Plasma enzyme measurements revealed that MANF deficiency alone did not affect circulating amylase or lipase activities (Fig. 8C). Caerulein treatment, however, induced approximately a threefold increase of amylase and lipase activities in the plasma. The combination of MANF deficiency and caerulein further elevated plasma lipase activity, but not amylase. Alcohol alone had no significant impact on amylase and lipase activities in the plasma, but when combined with MANF deficiency, it led to a pronounced increase in plasma lipase, while amylase remained unaffected.

**Figure 8.**
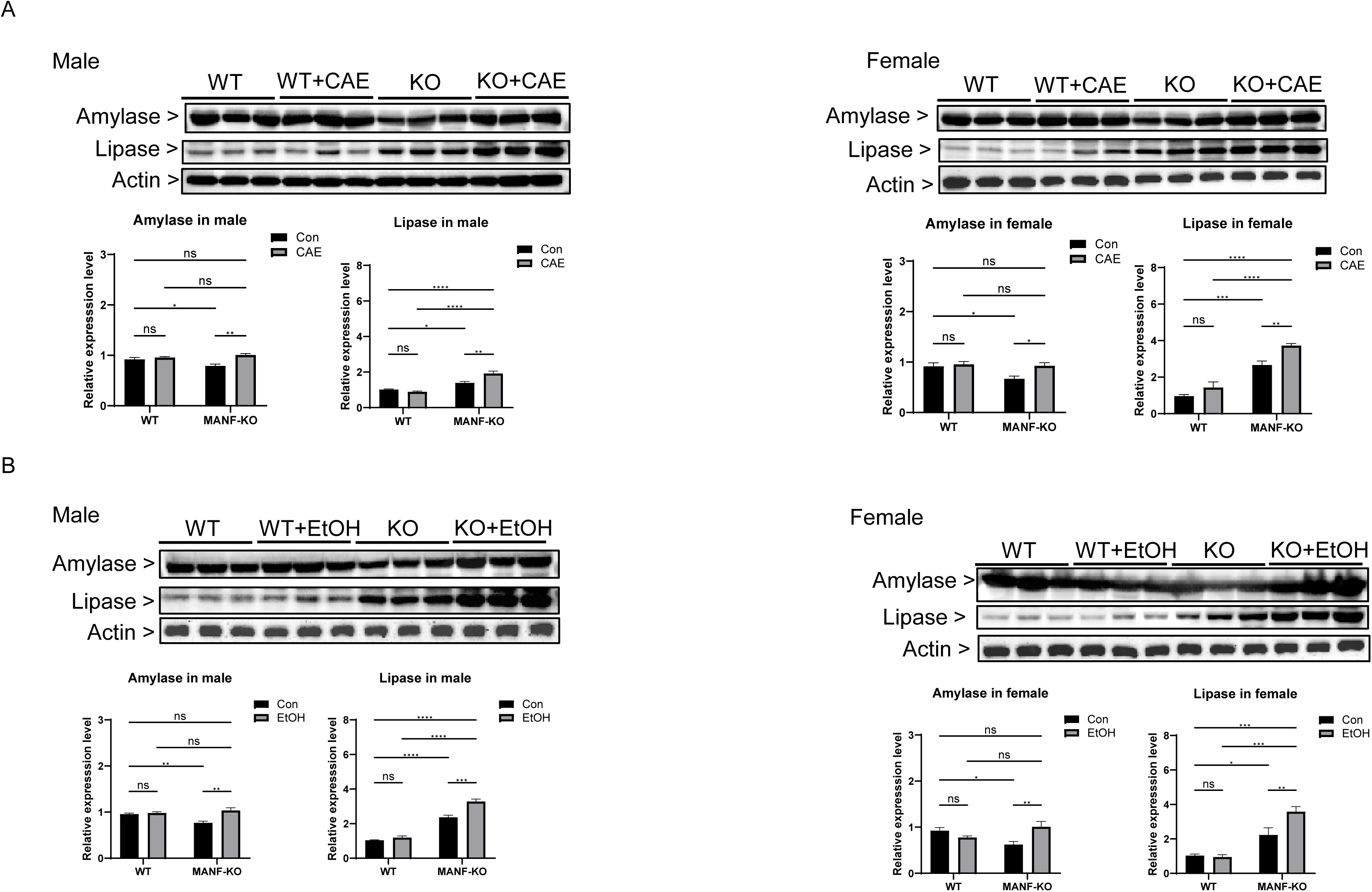
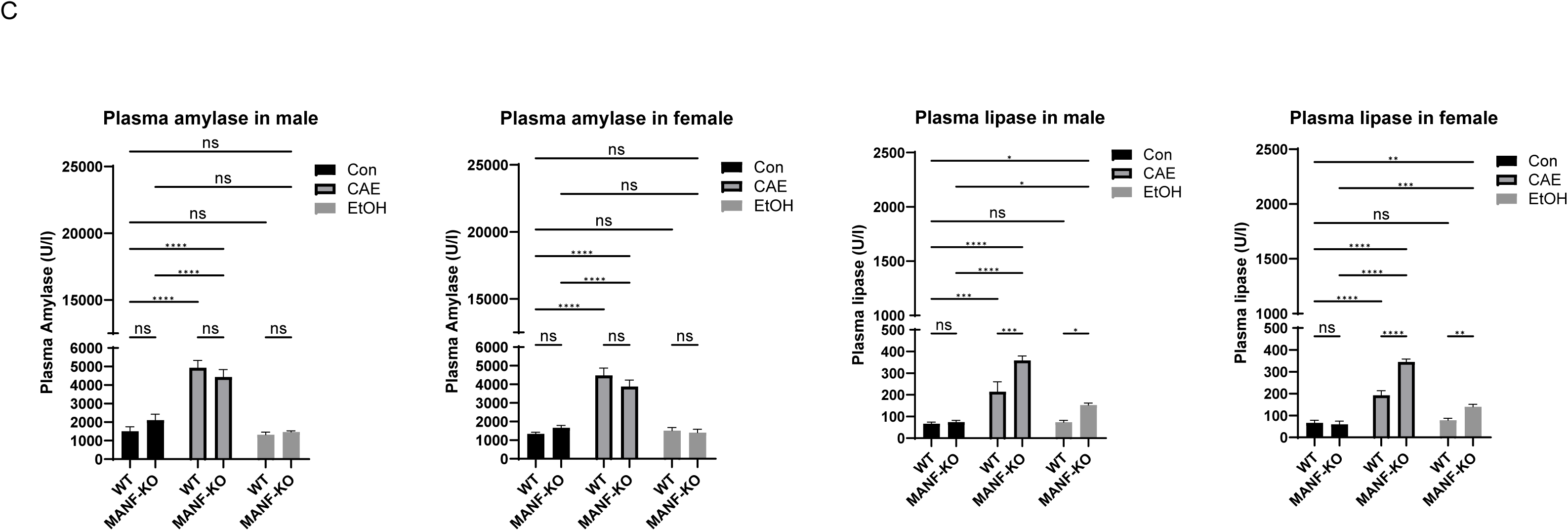
Effects of MANF deficiency on digestive enzymes in the pancreas of caerulein- and alcohol-treated mice. **A**. Representative IB image showing the expression of pancreatic amylase and lipase in male (left) and female (right) mice following caerulein treatment. **B**. Representative IB image showing the expression of amylase and lipase in male (left) and female (right) mice following alcohol exposure. n = 3 per group. **C**. The enzymatic activity of plasma amylase (left panels) and lipase (right panels) in male and female mice were measured by an enzymatic activity assay kit as described in Materials and Methods following caerulein or alcohol treatment. n = 4 per group. Data are presented as mean ± SEM. Statistical analysis was performed using two-way ANOVA with Tukey’s post hoc test. Significance is indicated as follows: *p < 0.05, **p < 0.01, ***p < 0.001, ****p < 0.0001; ns, not significant (p > 0.05).

### Effects of MANF deficiency on cell proliferation in the pancreas of caerulein- or alcohol-induced AP models

AP induces inflammation and acinar cell injury, which in turn triggers a compensatory regenerative response characterized by increased cellular proliferation ^31, 32^. To assess this proliferative activity, we performed IHC staining for Ki67, a widely used marker of cell proliferation, on pancreatic tissue sections (Fig. 9). Quantification of Ki67-positive cells across treatment groups showed that only caerulein treatment increased Ki67 expression in male mice. (Fig. 9B). Nor did MANF deletion by itself alter cell proliferation. However, the combination of MANF deficiency with either caerulein or alcohol treatment led to a marked and synergistic increase in Ki67-positive cells (Figs. 9A and 9B). These proliferating cells were primarily localized to the acinar and ductal regions, suggesting an enhanced regenerative response in these compartments under conditions of combined injury and MANF deficiency.

**Figure 9.**
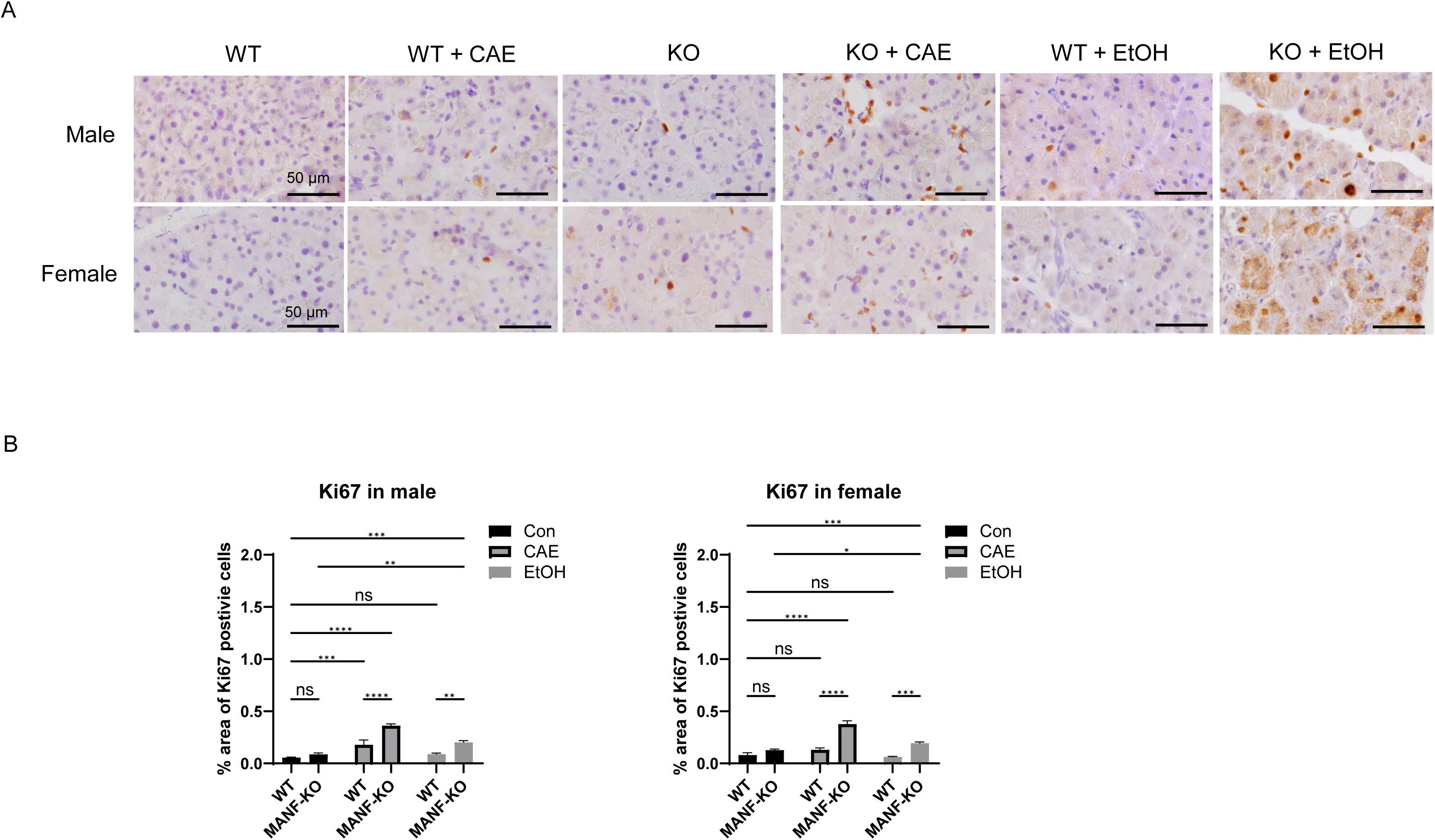
Effects of MANF deficiency on cell proliferation in the pancreas of caerulein- and alcohol-treated mice. **A**. Representative IHC images of Ki67 staining (magnification of 40X) in male and female WT and MANF-KO mice. **B**. Quantification of Ki67-positive cells from five randomly selected fields in the pancreas per mouse. n = 4 mice per group. Data are presented as mean ± SEM and analyzed by two-way ANOVA with Tukey’s post hoc test. *p < 0.05; **p < 0.01; ***p < 0.001, ****p < 0.0001; ns = not significant (p > 0.05).

## Discussion

To investigate genetic factors that may contribute to the pathogenesis of AP, we used two moderate experimental AP models, caerulein- and alcohol-induced pancreatic injury, to examine the role of MANF, an ER stress-inducible protein, in modulating the severity of pancreatic injury. The caerulein model mimics acinar cell hyperstimulation via excessive activation of cholecystokinin (CCK) receptors, leading to dysregulated enzyme secretion, ER stress, and activation of pro-inflammatory signaling pathways ^24, 33^. In contrast, alcohol-induced AP involves a more complex pathophysiology, including the direct cytotoxic effects of ethanol and its metabolites such as fatty acid ethyl esters on acinar and ductal cells, along with oxidative stress and ER dysfunction ^34, 35^. Although mechanistically distinct, both models share key pathological features such as ER stress, inflammatory cell infiltration, and cell death, making them complementary platforms to investigate MANF’s function.

Using the Cre/loxP system, we generated pancreas-specific MANF knockout (*Pdx1^Cre/+^*:: *Manf^fl/fl^*) mice and found that MANF deficiency alone results in significant reductions in body weight, pancreatic mass, body fat, and femoral bone mineral density, particularly in female mice. MANF KO had little effect on the motor coordination and sociability, but modestly affected anxiety-like behaviors in a sex-specific manner. Male MANF-KO mice showed increased anxiety-like behavior, whereas females exhibited reduced anxiety as shown in the open field test. These findings suggest that MANF plays a broader role in metabolic homeostasis and sex-dependent behavioral responses beyond its pancreatic functions. Notably, pancreatic-specific MANF deletion induced elevated levels of cleaved caspase-3 even in the absence of external injury, indicating a basal pro-apoptotic state. This apoptotic tendency was further exacerbated following treatment with either caerulein or alcohol, demonstrating that MANF deficiency markedly intensified pancreatic damages. The concurrent induction of caspase-12 in these combined conditions supports the hypothesis that MANF loss sensitizes the pancreas through enhancement of ER stress-associated apoptotic pathways.

How about MANF deficiency on basal UPR of all three pathways? Mechanistically, our data indicate that MANF deficiency amplifies ER stress responses triggered by both caerulein and alcohol through both shared and distinct pathways. In the caerulein-induced AP model, the combination of MANF loss and caerulein administration led to a marked increase in GRP78, total eIF2α, and phosphorylated eIF2α (p-eIF2α), suggesting enhanced activation of the PERK-eIF2α branch of the UPR pathways. In contrast, in the alcohol-induced AP model, the combination of ethanol and MANF deficiency selectively upregulated p-IRE1, GRP78, and p-eIF2α, indicating a distinct pattern of UPR activation involving the IRE1 arm. These findings suggest that MANF modulates multiple UPR pathways in a context-dependent manner, with its regulatory role varying by the type of cellular insult. The overlapping induction of both apoptotic markers (cleaved caspase-3 and caspase-12) and ER stress markers (GRP78 and p-eIF2α) in both models supports a converging mechanism by which MANF deficiency exacerbates ER stress-mediated cell death. However, the differential responses-such as the selective upregulation of total eIF2α in the caerulein model and p-IRE1 in the alcohol model-underscore insult-specific patterns of UPR activation resulting from MANF deficiency.

We also examined the inflammatory response associated with MANF deletion. Loss of MANF alone led to elevated expression of the pro-inflammatory cytokines TNF-α and IL-6, indicating a baseline pro-inflammatory state even in the absence of external stressors. Upon exposure to either caerulein or alcohol, the expression of both cytokines increased further, suggesting a synergistic interaction between MANF deficiency and external insults in amplifying inflammation. Notably, the combination of MANF deficiency with both caerulein and alcohol treatments resulted in a marked sex-specific response in HMGB1 expression: a synergistic elevation in males and a reduction in females. This finding is particularly significant given that HMGB1 is a well-established pro-inflammatory mediator implicated in various diseases, including pancreatitis ^36^. Elevated HMGB1 levels have been correlated with increased disease severity in both experimental models and clinical studies of pancreatitis ^27, 37^. The male-specific increase and female-specific decrease in HMGB1 observed in our study suggest a heightened inflammatory sensitivity in males and a potentially protective mechanism in females, underscoring a sex-dependent differential immune response. Collectively, these findings indicate that while MANF deletion enhances inflammation in both sexes when combined with caerulein or alcohol, the effect is more pronounced in males, possibly driven by increased HMGB1 expression.

CD68 is a well-established marker for macrophages and is widely used to evaluate immune cell infiltration in various tissues, including the pancreas during inflammation ^38^. In this study, we observed that MANF deficiency led to increased baseline CD68 expression exclusively in male mice, suggesting a sex-specific predisposition to a pro-inflammatory state. In both caerulein- and ethanol-induced acute pancreatitis (AP) models, MANF deletion further enhanced CD68 expression in males, indicating an exacerbated inflammatory response. Interestingly, in the ethanol-induced AP model, CD68 expression was also upregulated in females when ethanol treatment was combined with MANF deficiency. This observation implies that CD68 may serve as a more sensitive indicator of macrophage activation in females under ethanol-induced stress. Together, these results support a modulatory role for MANF in regulating macrophage-mediated inflammation during pancreatitis in a context-dependent and sex-specific manner.

We also examined the effects of MANF deficiency on oxidative stress, given its well-established role in the initiation and progression of pancreatitis-associated tissue injury ^39, 40^. In our study, we found that MANF deficiency interacted with either caerulein or alcohol to elevate levels of oxidative stress markers, DNP and 4-HNE which are indicative of protein carbonylation and lipid peroxidation, respectively in male mice but not female mice. This sex-specific increase in oxidative stress suggests that male mice may be more vulnerable to oxidative damage when MANF function is compromised. These findings point to a potential sex-dependent susceptibility that may influence the pathophysiological responses to pancreatic injury in the absence of MANF. This observation is consistent with previous reports showing that male rodents exhibit lower pancreatic antioxidant defenses and higher levels of oxidative damage than females under high-fat diet conditions ^41^.

A hallmark of AP, both in clinical diagnosis and experimental models, is the elevation of pancreatic digestive enzymes, particularly amylase and lipase, with increased plasma levels reflecting acinar cell injury and inflammation ^42, 43^. An intriguing finding in this study is the differential regulation of these enzymes by MANF. Specifically, pancreatic lipase protein expression was significantly increased in both male and female MANF-deficient mice, and this upregulation was further exacerbated by caerulein or alcohol exposure. Correspondingly, plasma lipase activities were also elevated under these conditions, consistent with lipase’s role as a sensitive indicator of pancreatic damage. In contrast, amylase exhibited a markedly different pattern. Neither caerulein nor alcohol affected pancreatic amylase, which is consistent with that these models produced moderate pancreatic injury. Caerulein but not alcohol increased plasma amylase, suggesting that caerulein caused more pancreatic damages than alcohol. MANF deficiency reduced baseline pancreatic amylase levels in both sexes; however, caerulein and alcohol treatment reversed this reduction. MANF deficiency had no effect on plasma amylase. These findings suggest that the presence of MANF is necessary for maintaining physiological amylase expression. This differential regulation aligns with clinical observations in human AP, where lipase is generally considered a more specific and sensitive biomarker of pancreatic injury than amylase—particularly in alcohol-induced cases, where amylase levels often remain normal or only mildly elevated despite substantial tissue damage ^42, 44^.

The pancreas has a remarkable ability to regenerate after injury, particularly in the exocrine acinar cells, and this regeneration occurs rapidly following cell death and inflammation ^45, 46^. Only caerulein but not alcohol increased Ki67-positive cells in male mice, again indicating the moderate nature of these AP models. However, we observed a synergistic increase in Ki67-positive cells six hours after the first caerulein dose and four hours after the final alcohol dose in both male and female MANF KO mice, indicating that MANF deficiency exacerbated pancreatic injury. While caerulein- or alcohol-induced pancreatitis is known to stimulate acinar regeneration and cell proliferation, the changes depend on timing and severity of these treatments. For example, Wodziak et al. (2016) reported pancreatic regeneration one day after caerulein-induced acute pancreatitis in mice ^47^. Ren et al. (2016) showed that more severe alcohol exposure-ten days of alcohol gavage induced pancreatic injury, with regeneration occurring six hours after the final gavage ^27^. Rydzewska et al. (2001) used a chronic ethanol-containing diet followed by caerulein injection to induce acute pancreatitis in rats and observed increased BrdU labeling six hours after the first caerulein injection, indicating regeneration ^48^. The early induction of Ki67 in our caerulein- or alcohol-induced models likely reflects a “primed” state of pancreatic epithelial cells in the absence of MANF, driven by sustained inflammation (e.g., elevated IL-6 and TNF-α) and cell death (e.g., cleaved caspase-3). In this pro-inflammatory environment, acinar cells may already be poised to enter the cell cycle, with caerulein or alcohol acting as a second hit that rapidly triggers G1 entry and subsequent Ki67 expression. This observation supports a two-hit model of injury-induced regeneration, in which genetic predisposition or chronic stress sensitizes pancreatic cells to respond more rapidly and robustly to acute insults. These findings suggest that the absence of MANF may enhances inflammatory signaling which makes acinar cells susceptible to further injury induced by environmental insults.

In summary, our findings demonstrate that MANF deficiency sensitizes the pancreas to acute injury in both caerulein- and alcohol-induced models of moderate AP. This enhanced susceptibility is characterized by increased ER stress, cell death, inflammatory responses, macrophage infiltration, elevated levels of pancreatic digestive enzymes, and enhanced cellular proliferation. Collectively, these results support the role of MANF as a protective factor during the pathogenesis of AP. Future studies are necessary to determine whether the delivery of exogenous MANF may offer protection against AP.

## Abbreviations

AP: Acute pancreatitis
ATF6: Activating transcription factor 6
CAE: Caerulein
C-cas-3: Cleaved caspase-3
Cas12: Caspase 12
CD68: Cluster of Differentiation 68
CHOP: C/EBP homologous protein
DNP: Dinitrophenol
eIF2α: Eukaryotic Initiation factor 2 alpha
ERAD: ER-associated protein degradation
ER: Endoplasmic reticulum
H & E: Hematoxylin and Eosin staining
HMGB1: High mobility group box 1
HNE: 4-Hydroxynonenal
GRP78: Glucose-regulated protein 78
IL6: Interleukin-6
IRE1α: Inositol-requiring kinase 1 alpha
MANF: Mesencephalic astrocyte-derived neurotrophic factor
PERK: Protein kinase-like ER kinase
TNFα: Tumor necrosis factor-alpha
UPR: Unfolded protein response
XBP1: X-box binding protein 1

## Data availability statement

The original data presented in the study are available upon request to the corresponding authors.

## Ethics statement

The animal study was approved by the Institutional Animal Care and Use Committee (IACUC) of the University of Iowa. The study was conducted in accordance with the local legislation and institutional requirements.

## Acknowledgement

The authors gratefully acknowledge the Comparative Pathology Laboratory (CPL), a research core within the Department of Pathology at the University of Iowa, for their expert histology services and technical support.

---

Supplementary Figure 1. Behavioral assessment of male and female MANF-KO and WT littermate mice at 8 weeks of age. **A.** Open field test: total distance traveled on Day 1 (top left) and Day 2 (top right), and time spent in the center zone on Day 1 (bottom left) and Day 2 (bottom right). **B.** Elevated plus maze: percentage of entries into open arms (left) and time spent in open arms (right). **C.** Rotarod test: latency to fall from the rotating rod. **D.** Three-chamber social interaction test: time spent interacting with the novel mouse in the social cylinder (left) and time spent in the social chamber (right). Group sizes: males – WT (*n* = 9), MANF-KO (*n* = 8); females – WT (*n* = 14), MANF-KO (*n* = 8). Data are presented as mean ± SEM and analyzed by two-way ANOVA with Tukey’s post hoc test. *p < 0.05; **p < 0.01; ***p < 0.001, ****p < 0.0001; ns = not significant (p > 0.05).

## Reference

[1] Yadav D, Lowenfels AB: The epidemiology of pancreatitis and pancreatic cancer. Gastroenterology 2013, 144:1252–61.

[2] Sankaran SJ, Xiao AY, Wu LM, Windsor JA, Forsmark CE, Petrov MS: Frequency of progression from acute to chronic pancreatitis and risk factors: a meta-analysis. Gastroenterology 2015, 149:1490–500. e1.

[3] Ali UA, Issa Y, Hagenaars JC, Bakker OJ, van Goor H, Nieuwenhuijs VB, Bollen TL, van Ramshorst B, Witteman BJ, Brink MA: Risk of recurrent pancreatitis and progression to chronic pancreatitis after a first episode of acute pancreatitis. Clinical gastroenterology and hepatology 2016, 14:738–46.

[4] Hart PA, Bradley D, Conwell DL, Dungan K, Krishna SG, Wyne K, Bellin MD, Yadav D, Andersen DK, Serrano J: Diabetes following acute pancreatitis. The Lancet Gastroenterology & Hepatology 2021, 6:668–75.

[5] Li H, Wen W, Luo J: Targeting endoplasmic reticulum stress as an effective treatment for alcoholic pancreatitis. Biomedicines 2022, 10:108.

[6] Drake M, Dodwad S-JM, Davis J, Kao LS, Cao Y, Ko TC: Sex-related differences of acute and chronic pancreatitis in adults. Journal of clinical medicine 2021, 10:300.

[7] Li S, Gao L, Gong H, Cao L, Zhou J, Ke L, Liu Y, Tong Z, Li W: Recurrence rates and risk factors for recurrence after first episode of acute pancreatitis: A systematic review and meta-analysis. European Journal of Internal Medicine 2023, 116:72–81.

[8] Shen H-N, Wang W-C, Lu C-L, Li C-Y: Effects of gender on severity, management and outcome in acute biliary pancreatitis. PLoS One 2013, 8:e57504.

[9] Sharma S, Aburayyan K, Aziz M, Acharya A, Vohra I, Khan A, Haghbin H, Nehme C, Ghazaleh S, Weissman S: S0078 Gender Differences in Outcomes of Acute Pancreatitis in Hospitalized Patients: Results From Nationwide Analysis. Official journal of the American College of Gastroenterology| ACG 2020, 115:S38–S9.

[10] Murugan NJ, Voutsadakis IA: Proteasome regulators in pancreatic cancer. World journal of gastrointestinal oncology 2022, 14:38.

[11] Cooley MM, Thomas DDH, Deans K, Peng Y, Lugea A, Pandol SJ, Puglielli L, Groblewski GE: Deficient Endoplasmic Reticulum Acetyl-CoA Import in Pancreatic Acinar Cells Leads to Chronic Pancreatitis. Cellular and molecular gastroenterology and hepatology 2021, 11:725–38.

[12] Hetz C, Axten JM, Patterson JB: Pharmacological targeting of the unfolded protein response for disease intervention. Nature chemical biology 2019, 15:764–75.

[13] Fu Y, Lucas AL: Genetic Evaluation of Pancreatitis. Gastrointestinal endoscopy clinics of North America 2022, 32:27–43.

[14] Ren Y, Liu W, Zhang J, Bi J, Fan M, Lv Y, Wu Z, Zhang Y, Wu R: MFG-E8 Maintains Cellular Homeostasis by Suppressing Endoplasmic Reticulum Stress in Pancreatic Exocrine Acinar Cells. Frontiers in cell and developmental biology 2021, 9:803876.

[15] Kim Y, Park S-J, Chen YM: Mesencephalic astrocyte-derived neurotrophic factor (MANF), a new player in endoplasmic reticulum diseases: structure, biology, and therapeutic roles. Translational Research 2017, 188:1–9.

[16] Danilova T, Lindahl M: Emerging roles for mesencephalic astrocyte-derived neurotrophic factor (MANF) in pancreatic beta cells and diabetes. Frontiers in Physiology 2018, 9:1457.

[17] Danilova T, Galli E, Pakarinen E, Palm E, Lindholm P, Saarma M, Lindahl M: Mesencephalic astrocyte-derived neurotrophic factor (MANF) is highly expressed in mouse tissues with metabolic function. Frontiers in Endocrinology 2019, 10:765.

[18] Caldwell NJ, Li H, Bellizzi AM, Luo J: Altered MANF expression in pancreatic acinar and ductal cells in chronic alcoholic pancreatitis: a cross-sectional study. Biomedicines 2023, 11:434.

[19] Wu H, Li H, Wen W, Wang Y, Xu H, Xu M, Frank JA, Wei W, Luo J: MANF protects pancreatic acinar cells against alcohol-induced endoplasmic reticulum stress and cellular injury. Journal of Hepato-Biliary-Pancreatic Sciences 2021, 28:883–92.

[20] Lindahl M, Danilova T, Palm E, Lindholm P, Voikar V, Hakonen E, Ustinov J, Andressoo J-O, Harvey BK, Otonkoski T: MANF is indispensable for the proliferation and survival of pancreatic β cells. Cell reports 2014, 7:366–75.

[21] Danilova T, Belevich I, Li H, Palm E, Jokitalo E, Otonkoski T, Lindahl M: MANF is required for the postnatal expansion and maintenance of pancreatic β-cell mass in mice. Diabetes 2019, 68:66–80.

[22] Wang Y, Wen W, Li H, Clementino M, Xu H, Xu M, Ma M, Frank J, Luo J: MANF is neuroprotective against ethanol-induced neurodegeneration through ameliorating ER stress. Neurobiology of disease 2021, 148:105216.

[23] Hingorani SR, Petricoin EF, Maitra A, Rajapakse V, King C, Jacobetz MA, Ross S, Conrads TP, Veenstra TD, Hitt BA: Preinvasive and invasive ductal pancreatic cancer and its early detection in the mouse. Cancer cell 2003, 4:437–50.

[24] Ding S-P, Li J-C, Jin C: A mouse model of severe acute pancreatitis induced with caerulein and lipopolysaccharide. World journal of gastroenterology 2003, 9:584.

[25] Choi SB, Bae G-S, Jo I-J, Seo S-H, Kim D-G, Shin J-Y, Hong S-H, Choi B-M, Park S-H, Song H-J: Protective effects of lithospermum erythrorhizon against cerulein-induced acute pancreatitis. Pancreas 2015, 44:31–40.

[26] Wen W, Li H, Lauffer M, Hu D, Zhang Z, Lin H, Wang Y, Leidinger M, Luo J: Sex-specific effects of alcohol on neurobehavioral performance and endoplasmic reticulum stress: an analysis using neuron-specific MANF deficient mice. Frontiers in pharmacology 2024, 15:1407576.

[27] Ren Z, Wang X, Xu M, Yang F, Frank JA, Ke ZJ, Luo J: Binge ethanol exposure causes endoplasmic reticulum stress, oxidative stress and tissue injury in the pancreas. Oncotarget 2016, 7:54303–16.

[28] Ren Z, Yang F, Wang X, Wang Y, Xu M, Frank JA, Ke ZJ, Zhang Z, Shi X, Luo J: Chronic plus binge ethanol exposure causes more severe pancreatic injury and inflammation. Toxicology and applied pharmacology 2016, 308:11–9.

[29] Xu H, Liu D, Chen J, Li H, Xu M, Wen W, Frank JA, Grahame NJ, Zhu H, Luo J: Effects of chronic voluntary alcohol drinking on thiamine concentrations, endoplasmic reticulum stress, and oxidative stress in the brain of crossed high alcohol preferring mice. Neurotoxicity research 2019, 36:777–87.

[30] Offield MF, Jetton TL, Labosky PA, Ray M, Stein RW, Magnuson MA, Hogan BL, Wright CV: PDX-1 is required for pancreatic outgrowth and differentiation of the rostral duodenum. Development 1996, 122:983–95.

[31] Malagola E, Chen R, Bombardo M, Saponara E, Dentice M, Salvatore D, Reding T, Myers S, Hills AP, Graf R, Sonda S: Local hyperthyroidism promotes pancreatic acinar cell proliferation during acute pancreatitis. The Journal of pathology 2019, 248:217–29.

[32] Murtaugh LC, Keefe MD: Regeneration and repair of the exocrine pancreas. Annual review of physiology 2015, 77:229–49.

[33] Kim H: Cerulein pancreatitis: oxidative stress, inflammation, and apoptosis. Gut and liver 2008, 2:74–80.

[34] Barreto SG, Saccone GT: Alcohol-induced acute pancreatitis: the ‘critical mass’ concept. Medical hypotheses 2010, 75:73–6.

[35] Clemens DL, Schneider KJ, Arkfeld CK, Grode JR, Wells MA, Singh S: Alcoholic pancreatitis: New insights into the pathogenesis and treatment. World journal of gastrointestinal pathophysiology 2016, 7:48–58.

[36] Wagner M: A dangerous duo in adipose tissue: high-mobility group box 1 protein and macrophages. The Yale journal of biology and medicine 2014, 87:127.

[37] Shen X, Li W-Q: High-mobility group box 1 protein and its role in severe acute pancreatitis. World journal of gastroenterology: WJG 2015, 21:1424.

[38] Chistiakov DA, Killingsworth MC, Myasoedova VA, Orekhov AN, Bobryshev YV: CD68/macrosialin: not just a histochemical marker. Laboratory investigation 2017, 97:4–13.

[39] Abu-Zidan F, Bonham M, Windsor J: Severity of acute pancreatitis: a multivariate analysis of oxidative stress markers and modified Glasgow criteria. Journal of British Surgery 2000, 87:1019–23.

[40] Robles L, Vaziri ND, Ichii H: Role of oxidative stress in the pathogenesis of pancreatitis: effect of antioxidant therapy. Pancreatic disorders & therapy 2013, 3:112.

[41] Gómez-Pérez Y, Gianotti M, Lladó I, Proenza AM: Sex-dependent effects of high-fat-diet feeding on rat pancreas oxidative stress. Pancreas 2011, 40:682–8.

[42] Ismail OZ, Bhayana V: Lipase or amylase for the diagnosis of acute pancreatitis? Clinical biochemistry 2017, 50:1275–80.

[43] Yang X, Yao L, Fu X, Mukherjee R, Xia Q, Jakubowska MA, Ferdek PE, Huang W: Experimental Acute Pancreatitis Models: History, Current Status, and Role in Translational Research. Front Physiol 2020, 11:614591.

[44] Batra HS, Kumar A, Saha TK, Misra P, Ambade V: Comparative study of serum amylase and lipase in acute pancreatitis patients. Indian journal of clinical biochemistry : IJCB 2015, 30:230–3.

[45] DiMagliano MP: Pancreatic stress and regeneration. Gastroenterology 2011, 141:1155–8.

[46] Zhou Q, Melton DA: Pancreas regeneration. Nature 2018, 557:351–8.

[47] Wodziak D, Dong A, Basin MF, Lowe AW: Anterior Gradient 2 (AGR2) Induced Epidermal Growth Factor Receptor (EGFR) Signaling Is Essential for Murine Pancreatitis-Associated Tissue Regeneration. PLoS One 2016, 11:e0164968.

[48] Rydzewska G, Jurkowska G, Dziecioł J, Faszczewska A, Wróblewski E, Gabryelewicz A: Does chronic ethanol administration have influence on pancreatic regeneration in the course of caerulein induced acute pancreatitis in rats. Journal of physiology and pharmacology : an official journal of the Polish Physiological Society 2001, 52:835–49.

