## Supplementary figures and images for "Deficiency of Mesencephalic Astrocyte-derived Neurotrophic Factor Aggravates Acute Pancreatitis in Mice"

### Supplementary Figure 1

Supplementary Figure 1

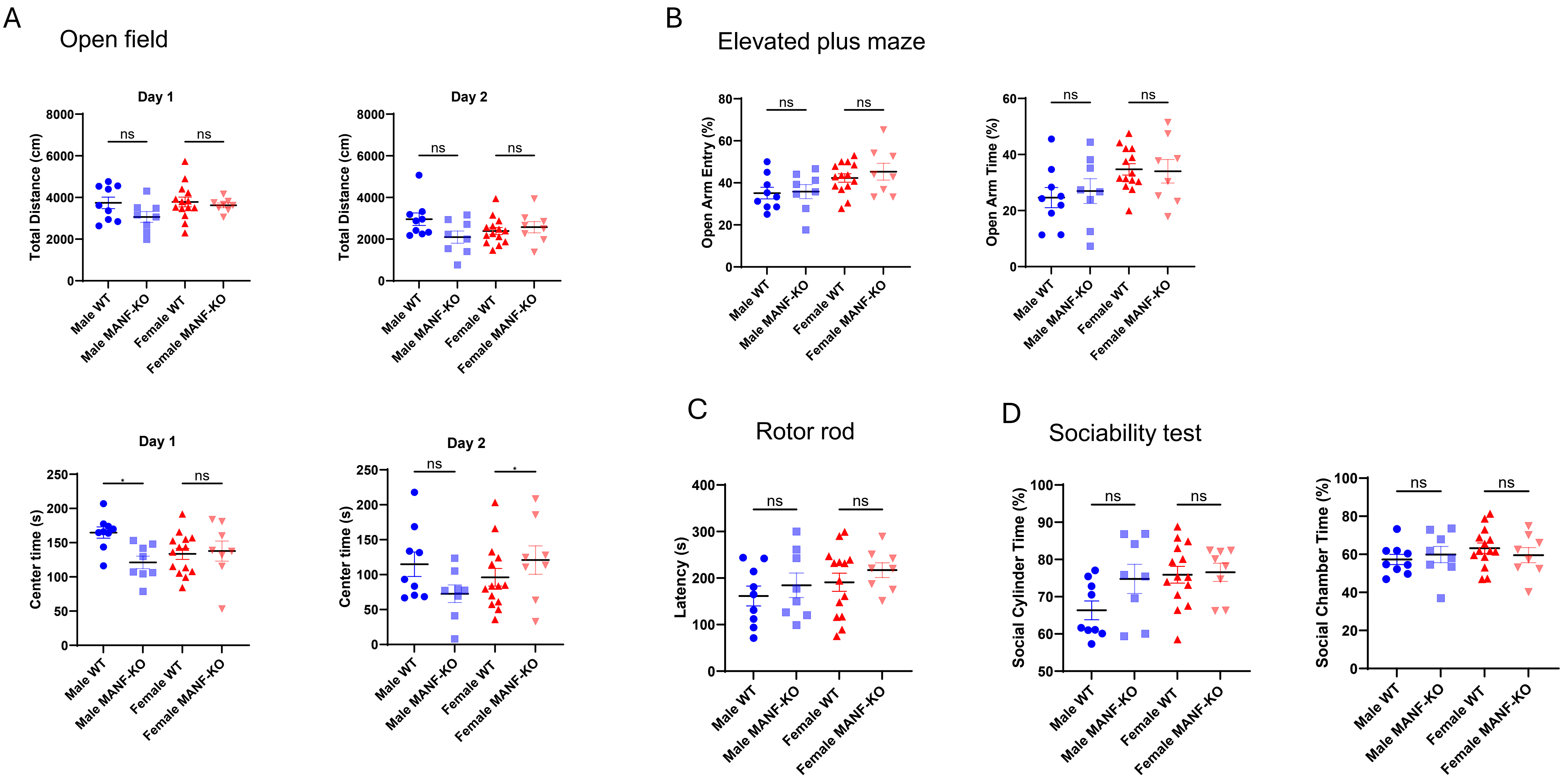
